# Tuning of Task-Relevant Stiffness in Multiple Directions

**DOI:** 10.1101/2025.02.13.637671

**Authors:** Chenguang Zhang, Federico Tessari, James Hermus, Himanshu Akolkar, Neville Hogan, Andrew B. Schwartz

## Abstract

The dynamics of any mechanical system can be described in terms of forces and motions. The interaction of these terms is often captured in the metric of mechanical impedance – a generalization of stiffness – used to describe how a mechanical system resists the application of force. The ability to adaptively change impedance is advantageous when encountering variable environmental conditions. Changing impedance in robotic systems is limited, whereas humans can rapidly and elegantly adapt the impedance of their limbs, especially during initial contact with objects. This is especially true for movements of our arms and hands. Multiple studies have examined the arm’s response to perturbation with the idea of impedance as a *reactive* component. In this study, we investigate the ability of humans to *predictively* set their arm impedance in a contact breaking task to perform fast movements to target positions in different directions. Our findings show that subjects (n=20) predictively co-activate antagonist muscles to primarily adjust one component of the arm’s impedance – stiffness – to match different task constraints before the movement begins irrespective of movement direction. Interestingly, the subjects’ performance was limited by the task-dependent stiffness rather than the required force and they tended to generate minimal stiffness to perform the task. With this robust strategy, task success is optimized at the expense of energy efficiency. This type of control is essential for the uniquely human ability to interact with objects.

## I. Introduction

The modulation and control of mechanical impedance^1^ is useful for describing the control of interaction in modern mechatronic and robotic systems (Hogan, 1980, 1985a). However, its importance in handling the transition between object interaction and free motion has received less consideration. Upon making or breaking contact, interaction forces change instantaneously, precluding or strongly limiting the continuous feedback control used in most robotic applications. Making or breaking contact not only causes an abrupt change in force that may induce bouncing, vibration, and instability, but also fundamentally changes the configuration of the control system. This is a fundamental challenge of robotics (Paul, 1987; Whitney, 1977).

In contrast, humans show a remarkable facility to adapt their motor control to physical interaction, and indeed take advantage of it. We use intrinsic mechanical compliance (more generally, mechanical impedance) as a key factor in the management of physical interaction (Kennedy & Schwartz, 2019a, 2019b). Adjustable mechanical impedance can minimize the need for rapid, continuous, and precise feedback, handles instantaneous force changes, and even incorporates initial contact as part of a control strategy, for instance, when the hand conforms passively to an object. For this type of interaction, two factors play a critical role: predictive tuning of the end-effector impedance and tuning of perturbation-induced reflexes. Our work focuses on the first aspect. The biological means to change intrinsic mechanical impedance is determined by skeletal configuration and the simultaneous activity of multiple muscles (Bennett et al., 1992; Franklin et al., 2003a; Hogan, 1985b; Perreault et al., 2001; Tee et al., 2004).

Cannon et al. (Cannon & Zahalak, 1982) showed that despite the neurophysiological complexity needed to control the mechanics of object interaction, the forces and resulting displacement of single joints are well described by a linear second-order dynamical model. The parameters of this model -- resistance (force) observed for variation of acceleration (mass), velocity (damping), and position (stiffness)-- competently describe the joint’s mechanical impedance. Impedance control was proposed as a theoretical framework, then rigorously extended to the neural, muscular, and skeletal systems to control multiple degrees of freedom posture and movement (Hogan, 1985a, 1985b). The work of Mussa-Ivaldi et al. (Mussa-Ivaldi et al., 1985) found evidence consistent with these theoretical considerations. Numerous studies have estimated limb mechanical impedance in different combinations of single-joint/multi-joint and static/motion conditions (Bennett et al., 1992; Franklin et al., 2003b; Gomi & Kawato, 1997; Krutky et al., 2013; Trumbower et al., 2009; Tsuji, 1997; van de Ruit et al., 2022)^2^. A consistent conclusion of these studies is that mechanical impedance at the hand is inherently highly configuration dependent.

A second factor contributing to the feed-forward cortical control of object interaction is the tuning of the stretch reflex that can also compensate for abrupt changes in muscle properties during task execution. The seminal work by Nichols and Houk (Nichols & Houk, 1973, 1976), and Crago et al. (Crago et al., 1976) highlighted how reflexes can compensate for variations in muscle stiffness. These results have been reconfirmed by more recent studies where subjects demonstrated the capability to tune their reflex activity during unexpected unloading tasks (Archambault et al., 2005; Reschechtko et al., 2019), or during unstable physical interaction tasks (Liaw et al., 2008).

In this work, we focus on predictive impedance control to generate movements during physical interaction. In this context, four prior studies are especially relevant to our study as they estimated impedance during forceful interaction tasks. Perreault et al. (Perreault et al., 2001) described how two factors, force level and limb configuration, influenced mechanical impedance, albeit, during static posture. Franklin et al. (Franklin et al., 2007) confirmed that the central nervous system is able to employ impedance control to overcome instability with the environment. Burdet et al. (Burdet et al., 2001) found that subjects changed their muscle activity in a way that increased limb stiffness to stabilize an unstable task but did not require the subjects to generate a specific target stiffness.

We are interested in the way interaction forces are controlled during the transition between breaking object contact and very fast unconstrained motion. This issue is similar to that posed by Lacquaniti et al. (Lacquaniti et al., 1993) who found that preparatory changes in muscular activity and impedance took place when catching a ball. In this type of task, object contact causes instantaneous changes in force and control must be predictive. To further study this type of control, we employed a ballistic-release paradigm (Kennedy & Schwartz, 2019b) in which subjects push and maintain a force in different directions against a handle which releases suddenly. The subjects then arrest the freely moving handle at a target location.

In the study reported here, we addressed (i) whether humans are capable of tuning task-specific stiffness – dictated by task demands – independently of the highly directional (configuration dependent) behavior of end-effector stiffness -- and (ii) what factors may limit the performance of interactive tasks. We asked subjects to perform a demanding planar ballistic-release task (left/right and forward/backward motions in a horizontal plane), without corrective movements. To perform this task, subjects could either employ very fast feedback control or they could set their arm impedance predictively to reach the target position. Subjects solved the task using the latter strategy.

We found that end-effector stiffness was invariant with respect to movement direction, depending only upon target force and displacement specifications. While previous work has demonstrated task-dependent changes in impedance (Burdet et al., 2001; Kennedy & Schwartz, 2019b; Lacquaniti et al., 1993; Perreault et al., 2001), our experimental design, which makes this task challenging, demonstrates that the factor limiting performance of this task is the ability to generate stiffness rather than force. In addition, subjects consistently completed the ballistic-release task with the minimum required stiffness, suggesting an effort-minimization strategy. Subjects predicted and adopted the appropriate stiffness consistently across different movement directions, showing that this control overcame the skeleto-muscular factors governing mechanical impedance. Pre-movement muscle co-activation was modulated with respect to task stiffness irrespective of the movement direction, suggesting that subjects tuned their mechanical impedance predictively. This learned strategy is efficient in that it minimizes the amount of information that needs to be transmitted during the movement, while being robust to context-dependent changes in the mechanics of the actuators.

## II. Methods

### A. Participants

20 healthy adults, ages 18 to 45, 13 males (height 160-188 cm and 50 – 113 kg weight) and 7 females (157.5 – 168 cm and 52.2 to 62.1 kg weight) participated in the experiment. All of them were right-handed; reported no neuro-muscular injury or disease; and consented in written form prior to their participation. The experiments conformed to a protocol reviewed and approved by the Institutional Review Board (IRB) at the University of Pittsburgh. Among the 20 tested subjects, one subject was eliminated from the analysis because the required experimental protocol was not followed correctly. In addition, subject 14 reported difficulties in successfully completing the lowest stiffness condition in the backward movement direction. This subject consistently fell short of the target position and compensated by making small terminal position corrections. The data from this subject in those specific conditions were therefore removed from our analysis.

### B. Robotic control

A WAM arm robot (Barrett Technology, Cambridge, MA), a 4-degree-of-freedom (DoF) back-drivable motorized linkage that allows control of force and impedance with a control loop frequency of 500 Hz, was used as the experimental platform. The robotic arm was configured to utilize its lowest-damping joint to minimize joint friction and other intrinsic non-linearities that could interfere with the task. The robot was set in its gravity-compensation mode to remove interactions due to gravitational forces.

The WAM was configured such that its endpoint moved primarily in a horizontal plane. Prior to release, the robot’s endpoint stiffness was maintained at 300 N/m on all three axes. This generated a point constraint at the manipulandum. Then, to implement the ballistic release, the stiffness was decreased abruptly to zero in the movement direction, while it was maintained at 300 N/m in the other axes. This produced a linear constraint on the motion of the manipulandum.

### C. Experimental setup

A handle was mounted on the WAM endpoint via a force transducer (Net F/T, ATI Industrial Automation, NC). An infra-red motion tracker (OPTOTRAK 3020, Northern Digital, Canada) recorded the handle’s 3D position at its tip. Other landmarks such as the subjects’ elbow and shoulder joints (identified by anatomical landmarks) were also recorded using the motion-tracking system. Subjects grasped the handle placed about 30 cm in front of their torso using their right hand as shown in Figure 1A. The gravitational load on the arm was canceled by a sling connected to the ceiling (Figure 1A) allowing the arm and forearm to move freely in a horizontal plane. Visual feedback was provided to the subject on a screen placed about 120 cm away from the subject’s eyes (Figure 1A). The feedback was composed of the planar position of the handle in the form of a filled circle, whereas the exerted force and the target position were presented as colored rectangular bars. To prevent excessive movement of the torso, subjects were harnessed to a wooden chair with a rigid back (harness not shown). A wrist brace was used to constrain wrist rotations.

**Figure 1.**
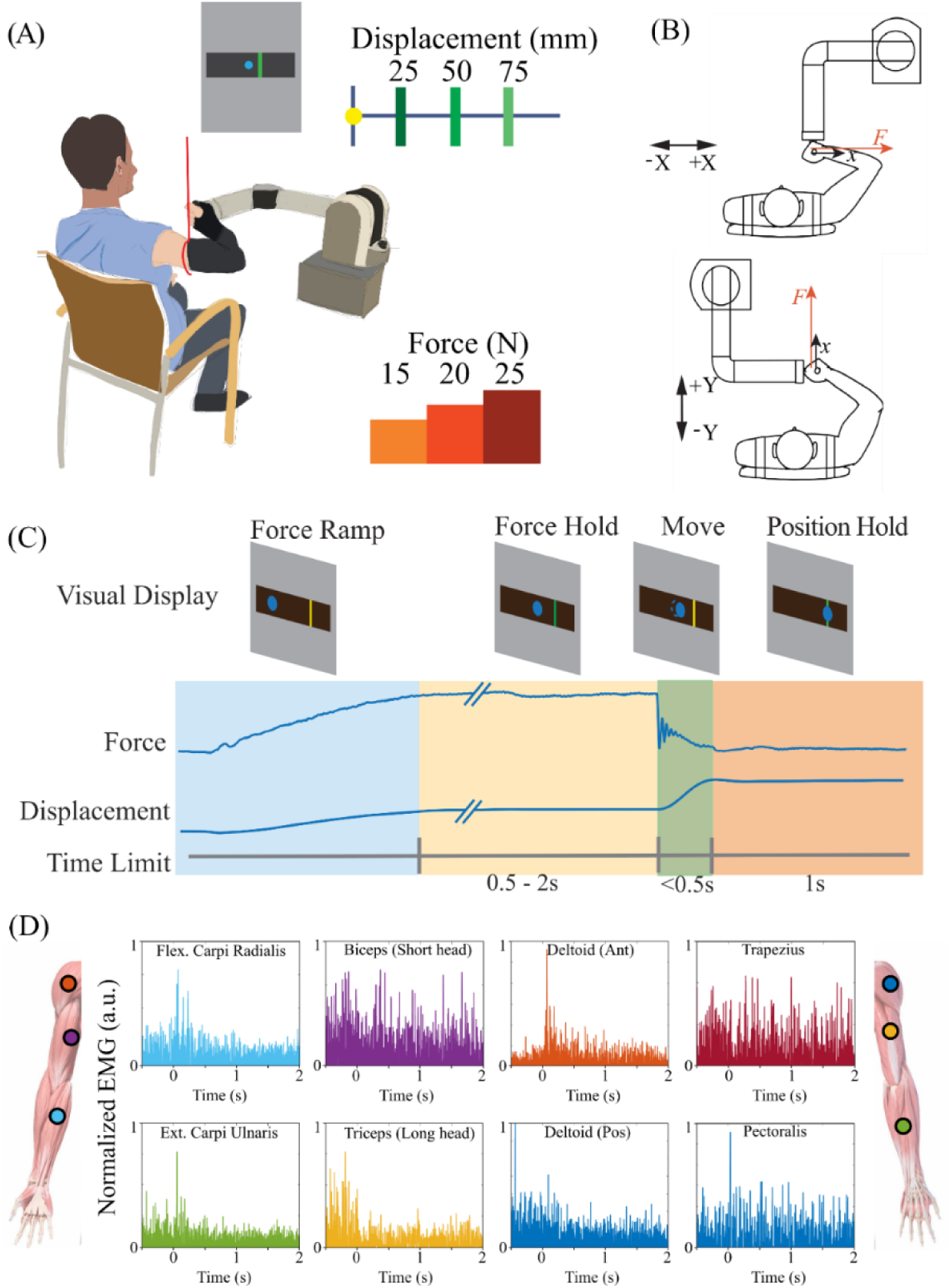
Experimental setup and design. **A**: Subjects sat in a chair and interacted with the WAM arm via the handle during the experiment. The subject’s arm was supported by a sling to avoid gravitational load and fatigue. A display screen provided feedback of the endpoint position and the force and position targets. **B**: WAM configurations for different directions. Subjects performed the task in four directions [+X/−X and +Y/−Y]. The subjects’ positions were changed for the task in different directions. **C:** Task-state diagram: A successful trial was composed of four consecutive phases: force ramp, force hold, ballistic move, and position hold. Subjects exerted and maintained a certain force against the handle for the first two phases. After a randomized duration (≥ 500ms) of force hold, the handle was suddenly released and began to move ballistically. Subjects were then required to maintain the target position (1 sec). The force exertion and ballistic movement occurred in the same direction. The position of the handle was depicted as a circular blue cursor; the target position and required interactive force were depicted as colored rectangular bars. The target bar during force hold was displayed as green when the correct force was exerted and was otherwise yellow or red when the exerted force was (respectively) lower or higher than the target range. During the movement phase, the bar turned green when the subject reached the required position target. **D:** Recorded, filtered (70-300 Hz) and rectified surface EMG from eight muscles. Signals were further smoothed (not shown here) for analysis (refer to Figures 9 and 10). Electrodes are shown in the approximate location of the targeted muscle groups.

Surface electromyographic (EMG) sensors were placed on eight upper-limb muscles. The signals were amplified (Octopus recording system, Bortec Biomedical, Canada) and recorded (Cerebus, Blackrock Neurotech, UT) at 2000 Hz. In post processing, the signals were first high-pass filtered at 10 Hz to remove any drift. The signals were band-pass filtered (70-300Hz) and rectified. The rectified signal, for each muscle, was then smoothed by applying a moving-average zero-lag filter with a fixed time window of *T*_*w*_ = 10 **m*s*. Finally, the individual EMG signals were normalized for each subject to the maximum EMG activity observed across all conditions and directions.

The task was performed for movements in four directions – Right (+X), Left (−X), Forward (+Y) and Back (−Y) as shown in Figure 1B. To avoid a possible confound due to robot behavior varying with direction, we fixed the robot configuration and rotated the subjects around it. Data collected during the task included the robot joint angles (robot encoders), the interactive force between the subject and the robot (force transducer), the subjects’ joint angles (infra-red motion tracker), and the handle position (infra-red motion tracker). A custom-built software system, RTMA, (Real-Time Messaging Architecture) (Velliste et al., 2008) was used to communicate between the different experiment modules to control the task and provide visual feedback to the subject.

### D. Trial paradigm

The experiment was designed to encourage subjects to preset their arm end-point impedance based on the force and displacement conditions prior to initialization of the movement. To complete this challenging task, it was necessary for subjects to learn, through trial-and-error, how to anticipate the mechanical aspects of the required movement. Learning was facilitated by employing a block design as explained in Section II E. A total of three forces [15N, 20N or 25N] (allowance ±2N) and three position targets [2.5cm, 5cm or 7.5cm] (allowance ±1cm) were selected such that the nine combinations of these parameters resulted in a wide range of expected stiffnesses from 200 N/m (15 N and 7.5 cm) up to 1000 N/m (25 N and 2.5 cm).

Subjects initiated a trial by pressing a button placed to one side of the handle. The button-press step allowed us: (i) to zero the force measurement to compensate for drift noise from the force sensor; and (ii) to set the robot to its home configuration for the upcoming trial. The button-press also helped subjects relax their arm, marking a distinct break between trials. After the button push, the actual task began. The task state (Figure 1C) was separated into four phases – force ramp, force hold, movement, and position hold.

Subjects were initially presented with a position target, which was displayed as a yellow vertical bar and the handle position as a blue cursor. Force feedback was presented by turning the bar green when the desired force during the force hold phase was achieved and yellow or red when force was (respectively) below or above the target force range.

For all analyzed directions [+X/−X, +Y/−Y], the direction of force generation and that of the movement were aligned. In other words, the ballistic release movement was performed in the same direction as the exerted force.

In the first two phases, force ramp and force hold, subjects grasped the handle and exerted force against the robot. When the target force was reached, the displayed bar turned green. The robot control was designed to guarantee that the hand position would be the same irrespective of the target force. Subjects had to maintain the exerted force within a range of ±2N for a random amount of time (0.5s to 2s) before it was removed abruptly, leading to a ballistic movement. A successful trial was accomplished if the subject stopped in the specified target zone and remained there for 1 second.

To ensure that subjects complied with the task specifications, performance was deemed a failure if: (i) the subject could not hold the force within the required range; (ii) the subject did not reach the position target range within 500 ms after release; or (iii) the subject made any corrective movement after stopping near the target. Corrective movements were detected as abrupt changes in velocity of the hand after the subject was in the position-hold phase of the trial. Minimizing corrective actions was intended to encourage subjects to predictively set the required end-effector arm impedance during the force hold phase. Subjects could move freely upon handle release with the stipulation they reached the target within 500 ms and maintained the final position.

To promote learning of the task specifications, the trials were performed in blocks, where each block consisted of repeated trials with the same force and position target conditions. Subjects had to successfully complete at least 9 trials to finish one block. The blocks were presented in a randomized order to prevent patterns of increasing or decreasing force or stiffness and to avoid subject fatigue. However, the sequence remained consistent across all subjects. Since we chose a combination of three force and three distance conditions, with each condition requiring nine trials, the subject performed a total of at least 81 successful trials in each direction. Nine blocks (three forces x three directions) were performed in each of the four directions: +X, −X, +Y, and −Y (Figure 1B). Lateral/medial movements were made in the “X” direction and “Y” denoted anterior/posterior motion. To familiarize the subjects with the task, we showed them a video clip describing the required task and allowed them to perform a tutorial block before beginning the experiment. The first condition block was repeated at the end of the experiment to mitigate any training effects. If the first and last blocks differed, the latter was used for data analysis. Learning effects were described by calculating the number of failed trials before a successful trial. If learning within a block took place, the number of failed trials before success should decrease with successive trials. For simplicity, we only used those trials in which the handle was released after the subject maintained the specified force level during the hold phase.

### E. Impedance parameter estimation

The arm end-effector behavior was modeled as a 2^nd^ order mechanical system i.e., a mass-spring-damper system as depicted in Figure 2.

**Figure 2.**
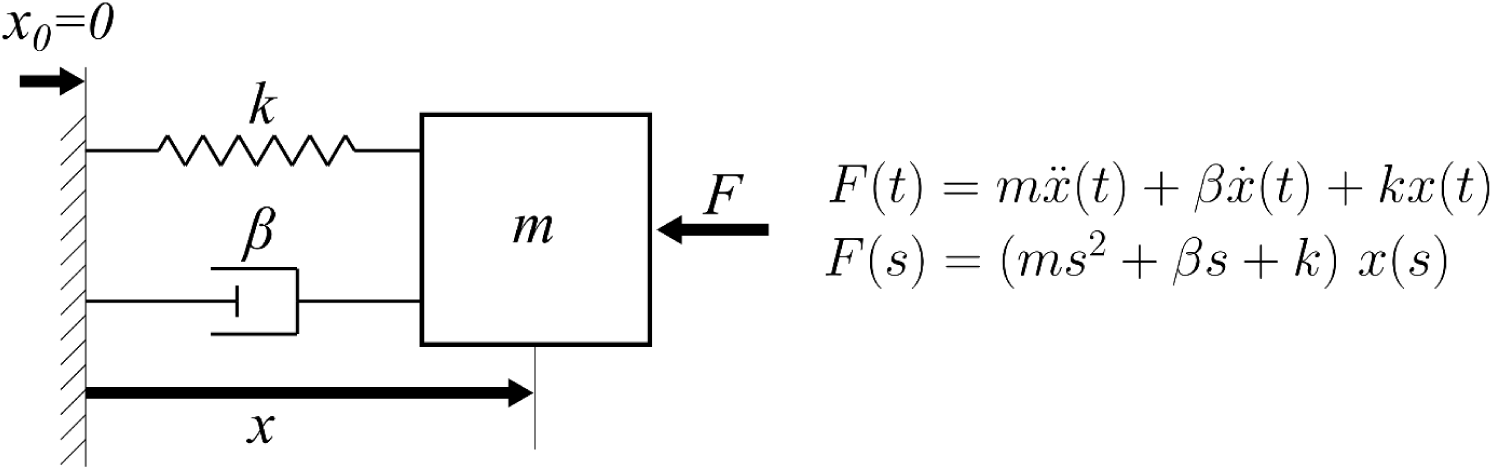
The 2nd-order dynamic model. The model describes arm behavior as a system with only 3 ideal components: a spring (*k*), damper (*β*), and mass (*m*). It also has a nominal position (*xx*_0_). The model input is force; the output is position.

The 2^nd^ order model used in our analysis contained stiffness (*k*), damping (*β*) and mass (*m*) coefficients corresponding, respectively, to the equivalent endpoint stiffness, damping and inertia apparent at the handle. Equation 1 presents the system transfer function.

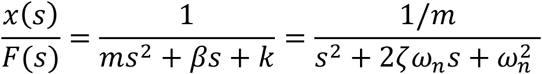

where *s* is the Laplace variable, 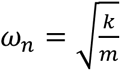 represents the system undamped natural frequency (rad/sec), and 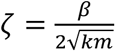 represents the system damping ratio (dimensionless).

We used the measured force and displacement at the handle to estimate the impedance parameters (mass, damping, and stiffness). These quantities were measured respectively by the force sensor mounted between the robot and the handle and by an optical marker placed on the tip of the handle and measured by the Optotrak system.

For each trial, the force and position recordings were used to perform parameter identification based on continuous-time system identification with the Instrumental Variable (IV) method (Garnier et al., 2010; Ljung, 2009; Young et al., 2007). MATLAB™’s system identification toolbox was used to perform the computation. The fit of the estimated model was computed as:

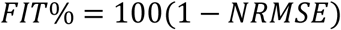

where NRMSE represents the normalized root mean squared error between the experimental displacement and that predicted by the identified second-order model.

### F. Statistical Analysis

The statistical analysis was performed using the SPSS statistical software package (SPSS Inc., Chicago, IL) with a significance level set to 5%. A statistical test was used to study the effect of direction on success rates. Two additional tests were performed to investigate the effects of the force-displacement conditions and of direction.

One-way, within-subject, repeated measures analysis of variance (ANOVA) was performed to examine the influence of the four different directions (+X, −X, +Y, and −Y) on the success rates of subjects. Force and displacement conditions were grouped for each subject. Post-hoc t-tests analyzing the main effects of the independent variable direction on the dependent measure were performed.

Linear regression was used to examine the influence of the estimated end-effector stiffness on the success rate in the four different directions (+X, −X, +Y, −Y).

The influence of target force and target displacement on the three dependent measures: stiffness, damping, and mass were examined with ANOVA. Separate analyses were conducted for the damping and mass dependent measures in four different directions: +X, −X, +Y, and −Y. For the stiffness dependent measure, a single ANOVA was performed by grouping the four different directions as random factors. In the case of significant interaction, post-hoc t-tests analyzing simple main effects of the independent variables force and displacement on the dependent measure were performed. ANOVA was also used to study how direction was related to stiffness, damping, and mass. Separate analyses were conducted for each of the 9 force-displacement conditions. Post-hoc t-tests analyzing main effects of the independent variable direction on the dependent measure were performed.

The p-values of post-hoc *t*-tests were adjusted using the Bonferroni procedure, where the original p-values were adjusted for the number of comparisons (*m*) by *p* = *p_*o*rig_* ⋅ *m* (standard method of Bonferroni correction in the SPSS software package adopted for statistical analysis). For the stiffness estimates, we also compared the mean stiffness – computed for each subject across trials and directions – with the expected stiffness at each target force and displacement condition. The expected stiffness was computed as the ratio between the target force *F_*tg*_* and the target displacement *d*_*tg*_.

The recorded and post-processed EMG signal (*⍺*) during the force-hold phase was averaged across trials for each condition, subject, and direction. We used a co-activation index (CAI) (Rudolph et al., 2000) as a measure of co-activation of the two pairs of agonist and antagonist muscles as per equation:

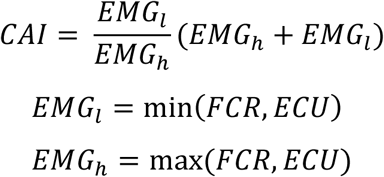

For each trial, EMG_l_ is the lowest of the average force-hold phase EMG recorded from the two muscles whereas EMG_h_ is the highest recorded EMG. The CAI is defined such that the simultaneous increase of both muscles will lead to a higher index, rather than individual muscle increase. The CAI was linearly regressed with the estimated task stiffness, force, and displacement (i.e., the conditions) for each subject and direction. The coefficient of determination *NN*^2^ was computed for each subject and direction.

## III. RESULTS

The following sections present the significant findings from our experiments. First, we explained the kinematics of the subjects using a simple a 2^nd^ order model (mass-spring-damper). The model fit the data well (Section III A) irrespective of whether the subject failed or succeeded in the trial. Next, we showed that subjects could learn the task-dependent impedance and that they were capable of adapting and learning different arm impedances (Section III B). The most critical parameter in describing the behavior was stiffness (Section III C) which was minimized by the subjects as they succeeded in the task (Section III D). Finally, we showed that subjects co-activated antagonist muscles before moving to set the stiffness of their arms predictively (Section III E).

### A. Ballistic release follows a 2^nd^ order model

Figure 3 shows an example of a recorded subject’s force and position data in one specific direction across the nine force-displacement conditions. Data were aligned at the time of handle release. Prior to release, the force was approximately constant and equal to the required target force ±2*NN*. At the time of release, the force decreased sharply towards zero and then changed in a complex manner as the arm moved smoothly toward the target. For higher force conditions, there was an overshoot in both force and direction about 250 ms after release. The subjects were allowed a maximum movement time of 500 ms.

**Figure 3.**
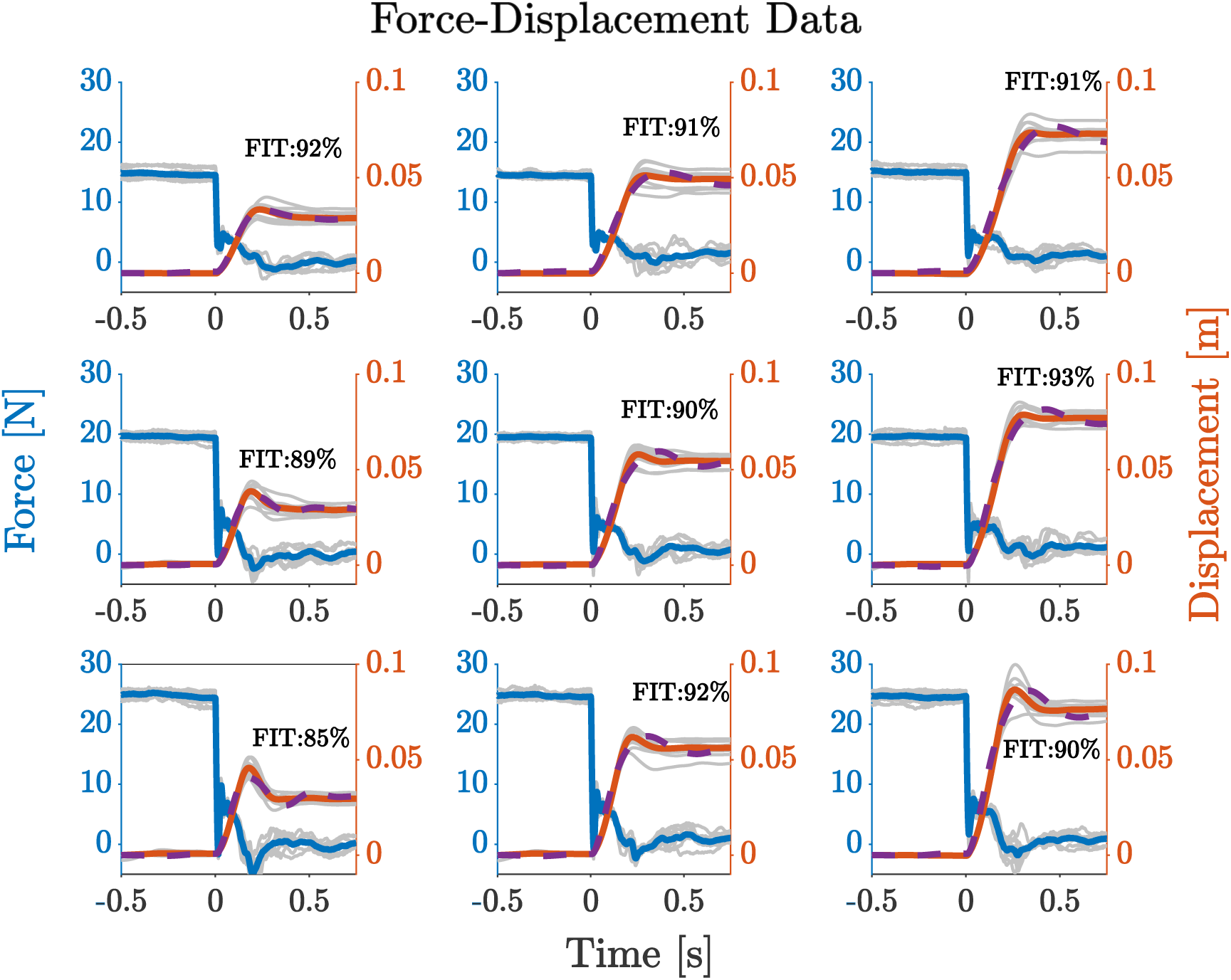
The force and displacement data for different conditions for subject 8 in direction +X. Target force (blue curves) conditions vary by row, while columns display different target displacement conditions (red curves). For each panel, the left y-axis shows the force (blue), and the right y-axis represents the displacement (red). The gray lines represent individual trials for each condition. The blue solid line shows the average force profile. The solid red line shows the average displacement. The dashed purple line shows the displacement predicted by the model. For every panel, we report the fitting performance of the model with respect to the experimental data.

The linear 2^nd^ order model had an excellent fit (FIT% > 85%) to the data in all trials and conditions (force and displacement), irrespective of failure or success (Supplementary Figure 4), even though several features of the underlying neurophysiology (e.g., reflexes) and mechanics (e.g., intrinsic friction of the robotic manipulandum) were omitted from the model. The effect of these omitted features can sometimes be seen in the force profiles of Figure 3 after release (*t* ≥ 0 *s*).

We found that the stiffness estimates from the second-order model were correlated with the task conditions (target force and target displacement). Mass estimates were roughly constant across conditions. This was expected, since, in a given direction, the overall mass of the robotic apparatus and the subject’s arm only negligibly changed due to a rotation of the end-effector mass ellipse. Consistent trends in the damping ratio were not observed across subjects. An in-depth analysis of the mass, damping and stiffness effects on movement distance are provided in the Supplementary Material (Section E, F and G).

### B. Subjects learnt to set the impedance to succeed the task

Subjects performed the task within blocks where for a given task condition, they had to succeed in 9 trials to complete the block. The performance across blocks within a single direction was described by a performance metric calculated as the number of failed trials before a successful trial was performed (Figure 4). The number of failures increased at the beginning of a new block (i.e. a new task condition) and decreased with successive trials within the block. This was especially evident for the more difficult cases such as those requiring high force and short displacements (marked by magenta arrows, Figure 4). Subjects had low failure rates in cases where the stiffness conditions were easier to achieve. The failure rate also dropped within a block, suggesting that the subjects were able to learn how to set the correct stiffness to complete the trial. These trends were observed in all directions. It should be noted that each trial began with a button push, disrupting the arm configuration used to hold the handle of the manipulandum at the beginning of each trial. The learned stiffness was retained even when the arm configuration changed transiently during button push. The first block (20 N, 50 mm) was repeated at the end of session and subjects again failed at the beginning of this repeated condition, suggesting that there was minimal carry-over of the learned behavior between blocks.

**Figure 4.**
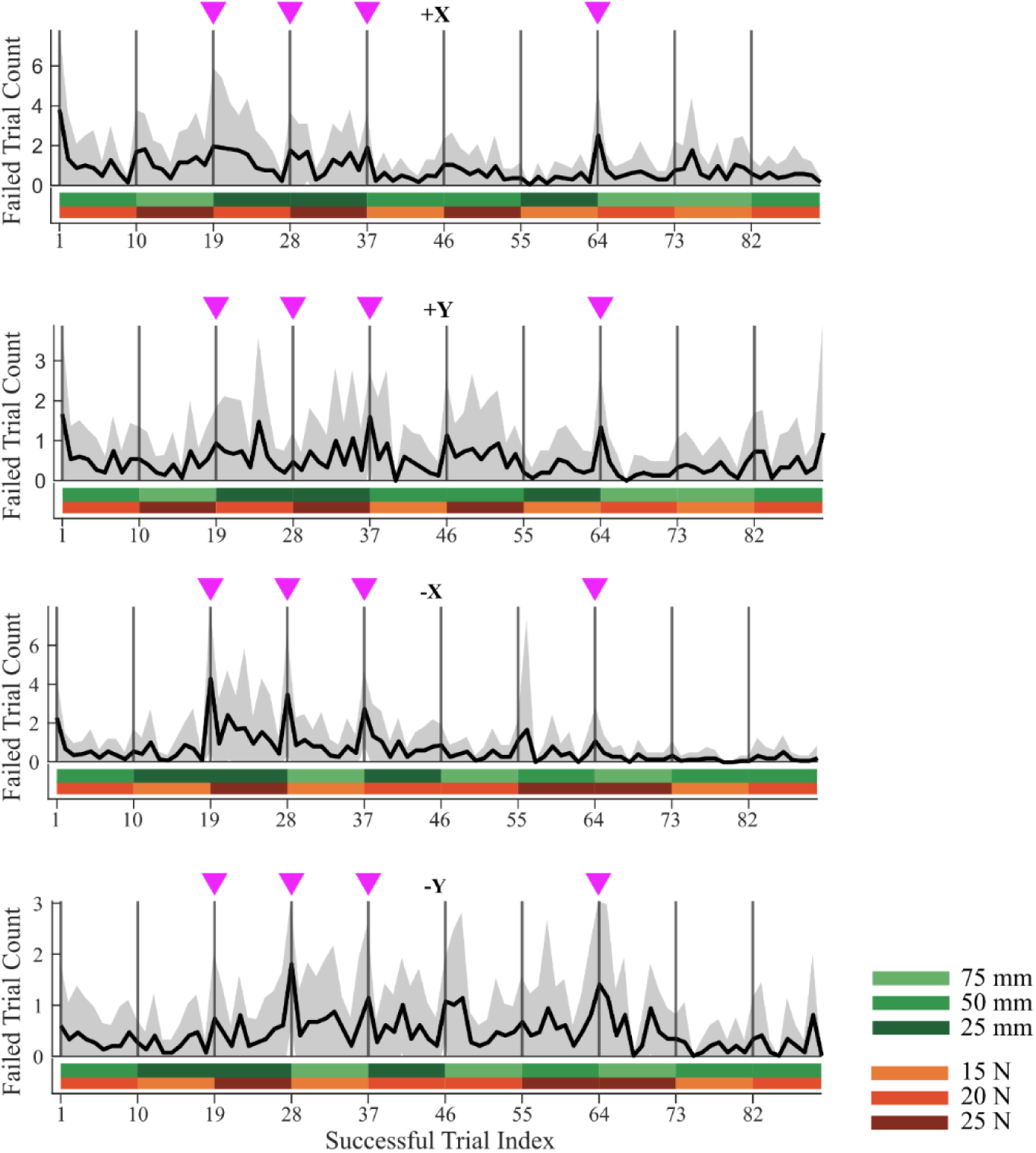
Failure rate over the entire session for the four directions averaged across all subjects. The vertical bars indicate the start of a new task condition. The subjects failed more trials when the condition was changed, especially in blocks highlighted by magenta arrows, as they required some trials before adapting to the new task conditions.

### C. Arm stiffness covaries with task condition

Results from the model fit the experimental data well for all subjects (FIT > 88%) and show that stiffness, damping, and mass for a given condition showed comparable values across subjects and directions (Figure 5 and Supplementary Figures 7,8). However, only stiffness was condition dependent. A sensitivity analysis showed that stiffness was the most powerful explanatory parameter as changes in its value caused large changes in the model fit (Supplementary Figure 2). Changes of ±30% from the estimated stiffness caused drops of the FIT of up to 70%. Equivalent damping and mass deviations from the estimated values showed a smaller effect on the FIT %. The averaged measured stiffness across subjects did not change with respect to movement direction in all 9 force-displacement conditions and was only affected by target displacement and target force—i.e., it was dictated by the task (Figure 5, bottom).

**Figure 5.**
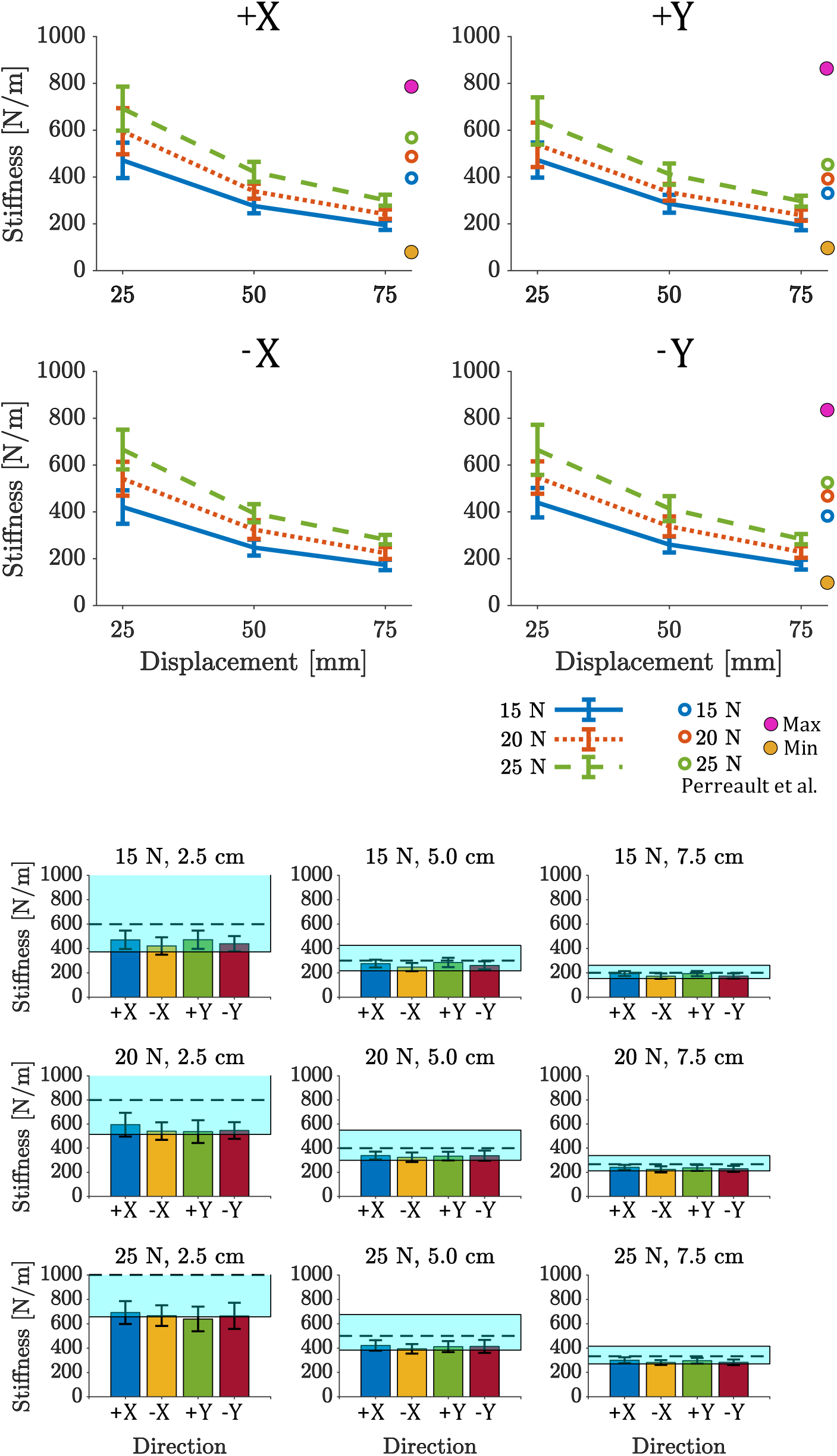
Arm stiffness variation across subjects. The top group of plots (4 plots) shows the stiffness change with respect to force and displacement in the four directions: +X (top left), +Y (top right), −X (bottom left), −Y (bottom right). For each panel, the nine different force-displacement conditions are presented: the x-axis shows the different target displacements, and the three colored lines represent the different target forces. The last column of vertical circles Y directions. The bottom group of plots (nine plots) shows the stiffness change in the different force-displacement conditions with respect to the four directions of motion. The light blue shaded areas represent the range of stiffness that subjects should reach to successfully complete the task, while the dashed line represents the expected stiffness. Error bars in both plots show one standard deviation.

Figure 6 shows the estimated stiffness for the successful and failed trials. We divided the data into low stiffness cases (less than 500 N/m) and high stiffness conditions (greater than 500 N/m). There was no difference between successful and failed trials for the mass and damping estimates. In contrast, subjects failed more often for the high stiffness conditions suggesting that they were unable to set their stiffness high enough in the failed trials. This suggests that stiffness may be the limiting factor for successful trials.

**Figure 6.**
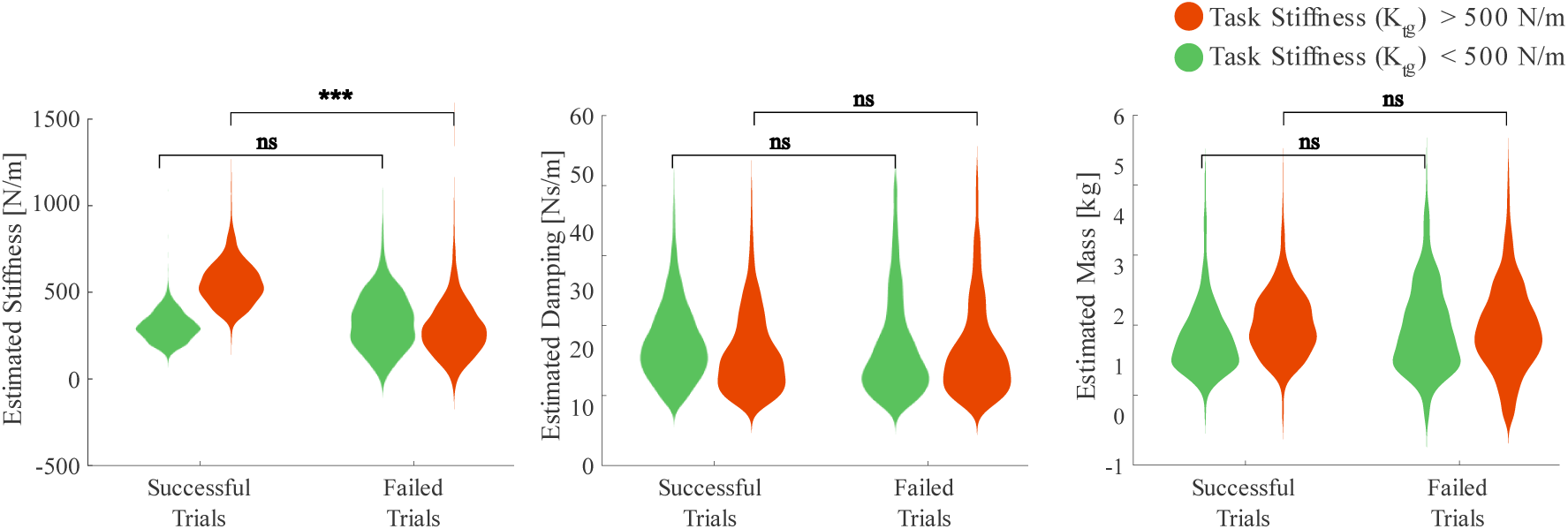
Parameter estimates for successful and failed trials. The data was divided into trials of high stiffness condition (nominal stiffness (Ktg) > 500 N/m, red) and low stiffness condition (Ktg < 500 N/m, green). For high stiffness conditions, the estimated stiffness was below the required stiffness when subjects failed the task. For low stiffness conditions the estimates for failed and successful trials were statistically similar. However, no difference was found in estimates of damping and mass between successful and failed trials suggesting stiffness being the critical parameter. (*** = p < 0.001)

ANOVA analyses indicated that force and displacement interacted across conditions and that both had significant effects on stiffness. Although there was a significant effect of direction on stiffness, it was not substantial (Supplementary Material).

### D. Subjects minimize end-point stiffness

Since we allowed some tolerance to the subjects for the required force (± 2*NN*) and the target position (± 1 *ccmm*), the equivalent stiffness is not a single value but rather a range as shown in Figure 7 (blue and green dashed lines). We found that the estimated stiffness for all subjects was always below the expected stiffness (specified force/specified target distance). The difference between the expected and estimated stiffness increased linearly with the expected stiffness. In fact, we observed that the average measured stiffnesses aligned well with the minimum stiffness that could be used as a single factor to complete the task.

**Figure 7.**
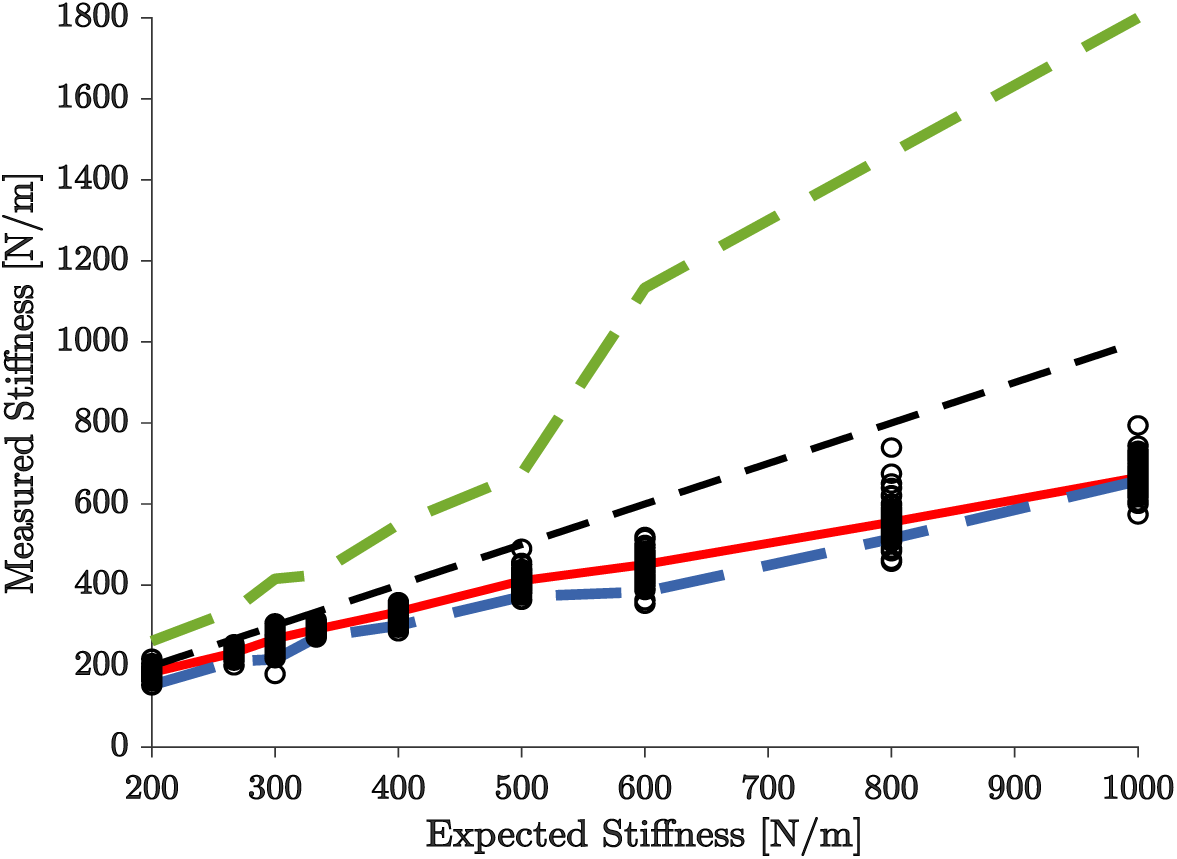
Comparison between the expected and experimentally measured stiffness. The expected stiffness is the ratio between the specified target force and the specified target displacement. Black circles represent the measured stiffness of each subject (across trials and directions), while the red line is the average measured stiffness across subjects. The dashed lines represent, from top to bottom, the maximum 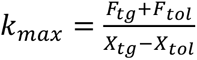 (green dashed), expected 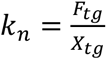 (black dashed), and minimum 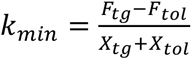 (blue dashed) stiffness using the assumption that this was factor dictated task success. *F_tg_*: Target force, *X_tg_*: Target displacement, *F_tol_*: Force tolerance = 2N; *X_tol_*: Displacement tolerance = 1cm.

Moreover, this was found to be true whether the subjects succeeded or failed the trial as shown in Figure 8. The figure shows the estimated stiffness for all subjects and all trials across the full session (shown here +X, +Y direction). The estimated stiffness (red and green lines) was always below the expected stiffness (blue bar) and at the lower end of the allowed range (blue shaded area) for all subjects. Similar behavior was observed in the −X −Y directions (Supplementary Figure 5). This suggests that subjects consistently set their stiffness near the minimum value needed to succeed in the task. It is important to note here that, in all the trials, the subjects could easily exert and maintain the desired force. Thus, it was the stiffness that became the limiting factor for the subjects to succeed in the task.

**Figure 8.**
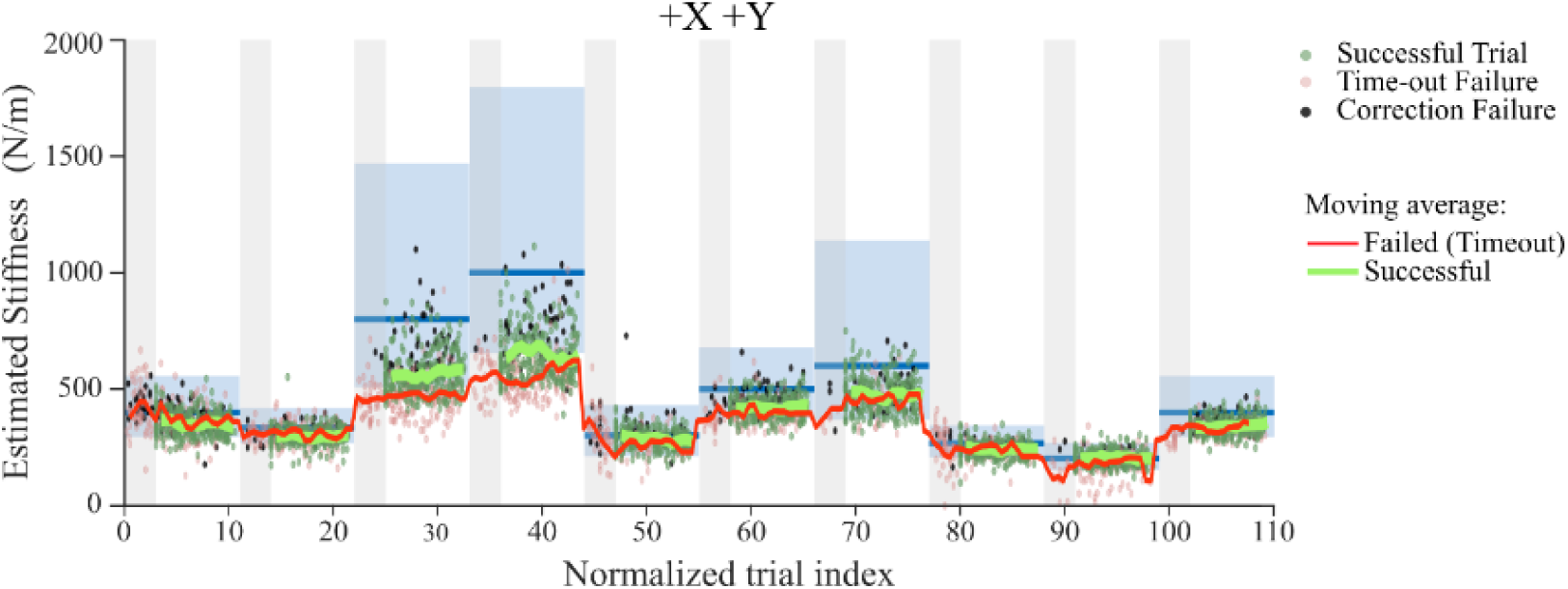
Stiffness estimates across sessions for all subjects in two directions. Gray sections show the failed trials between the last successful trial of one block and the first successful trial of the next. Dots represent the estimated stiffness for each trial where green represents a successful trial, red represents failure due to time-out and black dots show failed trials due to corrections during movement. The blue bars show expected stiffness 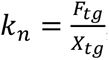, and the blue rectangle shows the range of stiffness based on the allowed force and displacement thresholds. The red line shows the average estimated stiffness for failed (time-out) trials whereas the green solid lines show the average estimated stiffness for successful trials.

### E. EMG shows predictive stiffness tuning by co-activation of muscles

Co-activation of antagonist muscles can increase the impedance of the limb. We recorded the EMG of eight arm muscles in our subjects and used a co-activation index (CAI (Rudolph et al., 2000)) as a metric of antagonist muscle pair co-activation. Although all muscles were differentially active across directions, only the wrist muscles (flexor carpi radialis and extensor carpi ulnaris) were consistently active in all directions (Supplementary Figure 13A). We used this pair as an example to show how EMG was related to the different task conditions (Figure 9), with the remaining muscle activations shown in Supplementary Figure 13B. In the pre-movement, force-hold portion of the task, the modulation of EMG activity was small, with a slight tendency to increase with higher stiffness conditions. The mean EMG for both flexor and extensor were well-correlated with stiffness (Figure 9B). The CAI for this pair of force-hold EMGs was calculated and found to be strongly correlated to the subject’s stiffness as it varied across task conditions (R^2^ = 0.81, Figure 9C). The coefficients of determination (*R*^2^) of the CAIs with respect to the estimated task stiffness, the magnitude of force during the force-hold, and the target displacement are plotted in Figure 9D. The CAI of these wrist muscles were well related to stiffness with a median *R*^2^ =0.64 and *IQ*_*R*_^2^ = [0.39 0.81]. The co-activation of the remaining muscle pairs was also well related (R^2^ = 0.62) but only in the directions in which they were active (Supplementary Figure 13B).

**Figure 9.**
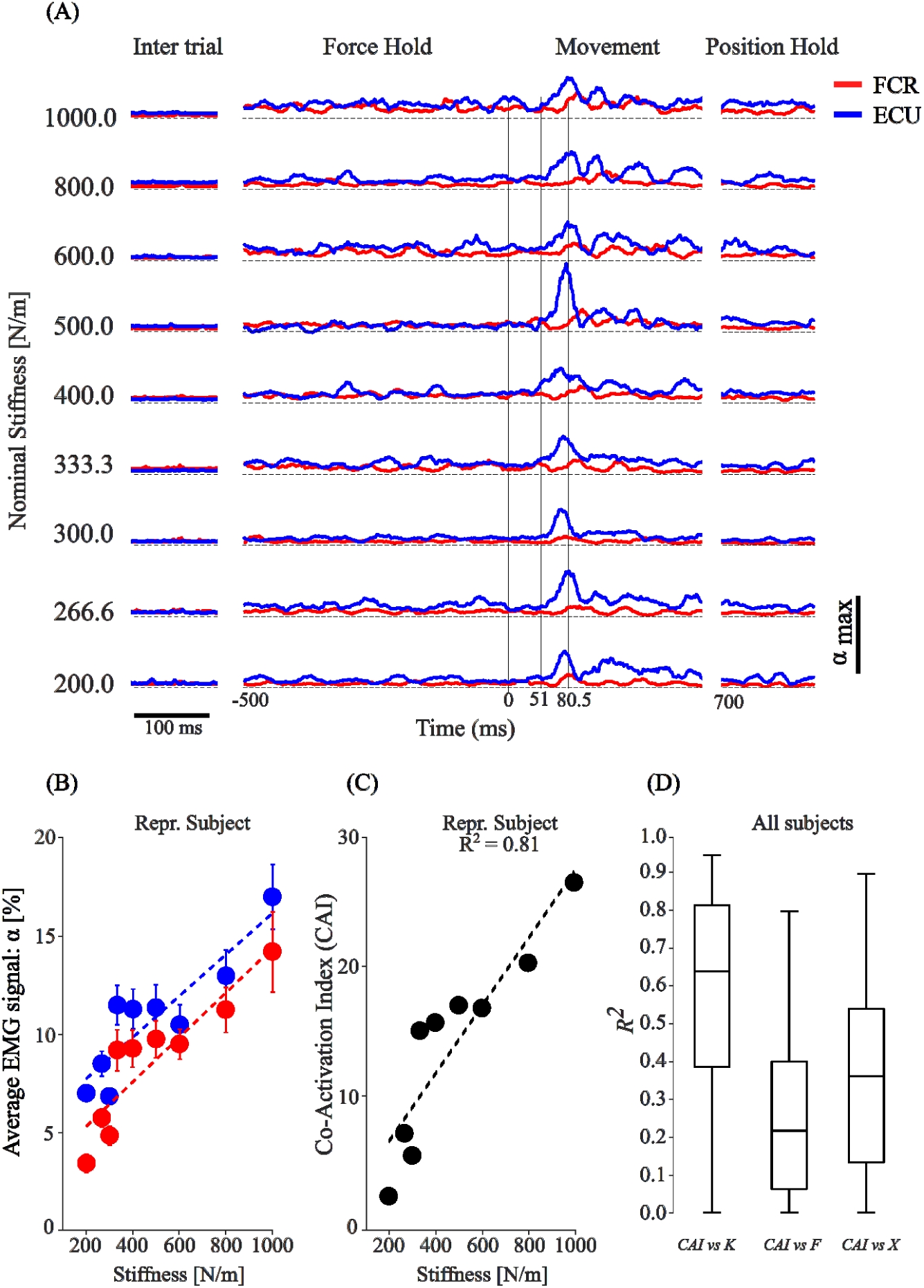
Task-related EMG. **A**. EMG recorded from FCR and ECU in a single subject moving in the −Y direction for the 9 different stiffness conditions. Each signal was normalized by its maximum value across all recordings (α_max_). The EMG of both muscles was fairly constant during the hold phase. Both increased with task stiffness compared to the intertrial interval. There was noticeable modulation of the ECU starting 30-50 ms after the handle was released (Time = 0) and peaking approximately 30 ms later. In the terminal hold phase, the EMG was again constant and similar to that during the pre-release hold period. **B**. EMG averaged across trials for different task stiffnesses during the hold phase. The EMG from both muscles increased with stiffness. **C**. A co-activation index (CAI) between the two muscles was calculated for each task condition and was highly correlated with the estimated stiffness. **D**. Across all subjects, the CAI was better related to stiffness (R^2^ = 0.64) than to force (R^2^ = 0.22) or displacement (R^2^ = 0.36).

Subjects changed their wrist muscle co-activation in a task-dependent manner to anticipate the motion of the manipulandum when it was released at the end of the position-hold portion of the task. This change can be interpreted as a task-dependent variation in wrist stiffness. After release, the handle moved rapidly, stretching and shortening muscles throughout the arm. Stretch-related reflexes are evident in the EMG activity. This reflex-related activity in the wrist extensor began approximately 50 ms after release and peaked about 30 ms later (Figure 9A). The reflex activity is further detailed in Figure 10. The wrist extensor EMG increased during the reflex but the response in the flexor did not (Figure 10A). This difference is emphasized in the mean EMG plots of Figure 10B. The wrist EMG of both the flexor and extensor muscles increased but not as strongly as in Figure 9B. Stiffness-related co-activation was also not strongly correlated to stiffness as evident in the reflex-response of this subject (R^2^ = 0.48, Figure 10C). Across subjects (Figure 10D), the correspondence between reflex-associated co-activation and stiffness was weaker than during the force hold period. This reduction in co-activation during the movement may be transient as the CAI-stiffness correspondence increased in the position-hold period to nearly the same values as the force-hold phase, across subjects (Supplementary Figure 14).

**Figure 10.**
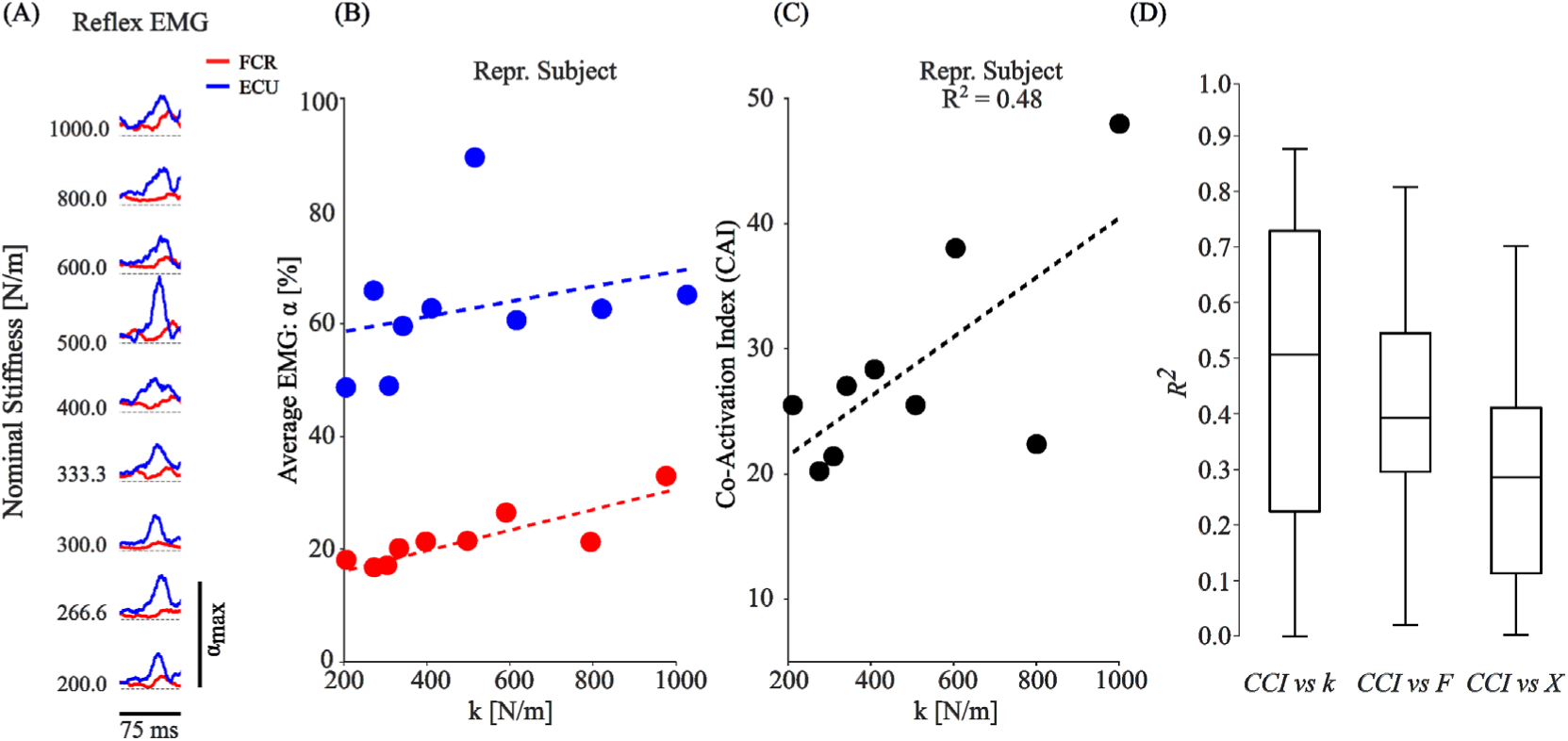
Reflex-associated EMG following release. A. FCR and ECU EMG across-stiffness conditions recorded from a single subject. The traces are aligned to their peak values following release. B. Mean EMG values from A versus task stiffness. C. CAI for the muscle pair versus task stiffness. D. Reflex-evoked CAI regressed against stiffness, force, and displacement for all subjects. Although the reflex-evoked CAI increased with increasing stiffness, this relation was weaker than during the pre-release hold phase.

## IV. Discussion

The human control of hand impedance/compliance far exceeds that of robots and even other primates (Orban et al., 2006; Stout & Chaminade, 2007; Walker, 2009), giving us an unparalleled ability to use tools such as a wrench to unscrew a stuck bolt. Our results show that, in challenging interaction conditions, subjects set their arm impedance even before the movement occurs as a strategy to achieve a goal within the constraints of both the task and the motor system. This strategy is robust to directional changes in the mechanics of the musculoskeletal system – which can be substantial – suggesting that anticipatory impedance setting may be a fundamental aspect of human motor control. Such a strategy may be employed in situations where neural sensory feedback would not allow for sufficient control. Common examples include extending the leg and foot to make surface contact during locomotion (Lee et al., 2016). The “roll-on” event between heel strike and foot flat occupies about 100 ms yet the neural transmission delay between peripheral sensing and spinally mediated mechanical responses is about 70 ms, while supra-spinally mediated responses may occur much later, suggesting that impedance is pre-set in anticipation of contact. Reaching to grasp an object provides another example. Conforming the fingers to exactly match the object shape (which may be poorly known i.e., when based on partially occluded visual data) is not necessary if the compliance of the hand and fingers is exploited (Lemerle et al., 2021). The idea that compliance could be simply controlled by setting a balance between pairs of opposing spring-like muscles was the essence of the “equilibrium-point hypothesis” proposed almost sixty years ago (Feldman, 1966). The general concept of predictive impedance setting is incorporated into the theory of “impedance control” (Hogan, 1984, 1985a).

Our results are consistent with a study using a similar paradigm in which perturbations were applied as subjects exerted force in two directions (Perreault et al., 2001). That study found a range of stiffness comparable to ours (see circles on the side of the plots in Figure 5, top panel). However, their study did not require subjects to anticipate an upcoming movement. The behavior we observed is consistent with the original equilibrium point hypothesis in which a pattern of agonist and antagonist muscle activation acts as a pair of opposing springs to set a balance or equilibrium position before movement begins (Feldman, 1966, 1986). Once moving, the limb will subsequently come to rest at the equilibrium point. While details of the movement trajectory could emerge from peripheral neuro-mechanics (Polit & Bizzi, 1978, 1979), subsequent research demonstrated that this was not a complete description of the neural control of reaching (Hinder & Milner, 2003; Lackner & Dizio, 1994). Nevertheless, the “final position control” version of the equilibrium point theories provides the most parsimonious account of our observations. Although we cannot rule out a pre-determined time course of neural activity based on some form of internal model of limb dynamics, the uncertain moment of release would make it difficult to time this neural trajectory appropriately and this may be a “good enough” form of control (Graziano & Gross, 1998), deployed when more sophisticated control strategies are unnecessary. This conclusion is supported by a recent study (Takagi et al., 2022), which used ballistic release in different directions to show that, when subjects generated different stiffness by changing grip force, their movements terminated at an equilibrium point in open-loop conditions. Future models of human motor control should be able to account for these characteristics of task-specific behavior (Hogan & Sternad, 2012; Schaal, 2006; Schaal et al., 2007).

### Task difficulty

The overall task was particularly challenging, showing an average success rate of about 64% ± 19% across different directions (Supplementary Figure 1). We used a block design where the task condition (specified force and displacement) was the same across repeated trials. This allowed subjects to learn how to set their arm impedance to compensate for the sudden handle release. Initial performance declined at the beginning of each block with a new condition, showing that negligible between-block learning took place, and then improved across repeated trials. In the higher stiffness conditions (short displacement and high forces), subjects had difficulty capturing the target and attributed this to a lack of strength. Indeed, Figure 6 shows that the subjects could not generate high enough stiffness in these cases, leading to failures. The damping and mass parameters remained unchanged between successful and failed trials suggesting that stiffness generation was the critical factor in the subjects failing the task. Interestingly, Figures 7 and 8 suggest that subjects consistently minimized the stiffness to complete the task, irrespective of whether they succeeded or not, even in case of low stiffness conditions. All subjects could generate three levels of force but struggled to reach high levels of stiffness (Figures 3, 6 and 7). This suggests that they were limited not by the amount of force required, but by the amount of stiffness.

### Variation of mass, damping, and stiffness

We found that a simplified 2 DoF (shoulder and elbow) arm model using arm inertial properties and the measured experimental force and displacement to calculate impedance (Cannon & Zahalak, 1982) described our data well (FIT% > 88%) whether the subjects succeeded or failed in the trials (Supplementary Figure 3). We used the force-displacement data from the entire task to accurately fit the stiffness. This suggests that despite the presence of higher order behavior such as stretch reflexes or friction in the robotic manipulandum, the stiffness remained roughly constant throughout the task.

The model showed that arm configuration had little bearing on the subjects’ presetting strategy (Supplementary Material). Subjects actively tuned their joint stiffness to meet the required forces and displacements specified by the task in each of the four directions. Furthermore, the stiffness estimates for each force-displacement combination were indistinguishable across directions. Although the intrinsic stiffness at the hand is known to be highly direction-dependent (Mussa-Ivaldi et al., 1985), subjects were able to pre-set their stiffness independently of movement direction.

Subjects tended to generate the minimum stiffness (below the nominal value) required to accomplish the task – the minimum target force over the maximum target displacement – and this trend persisted even when subjects failed. A smaller end-effector stiffness required a smaller joint stiffness, hence less muscle co-activation, and that would have reduced metabolic energy expenditure. This suggests that subjects tried to minimize the required muscle activity while meeting the task requirements and is consistent with the observation that in some circumstances (especially tasks requiring pushing), human neuro-muscular performance is not limited by force exertion but by stiffness production (Rancourt & Hogan, 2009).

### Pre-movement predictive tuning of stiffness

Pre-release EMG activity was well related to the estimated task stiffness across most subjects and directions. Specifically, the co-activation index (CAI) showed a good linear correlation with the estimated task stiffness whenever the muscles were found to be active (Figure 9C, D and Supplementary Figure 13). Release elicited a stretch reflex in the muscles of the arm and hand, which was evident in our EMG data. Although co-contraction also increased with stiffness during the reflex, the relation was weaker than the co-contraction-stiffness relation during the initial hold period (Figure 10). The latency of the spinal-mediated response is ∼40 ms and for the supraspinal contribution ∼75 ms (Cannon & Zahalak, 1982; Lewis et al., 2005) which is consistent with our result. Berret et al. (Berret et al., 2024), using a feed-forward model, showed that predictive co-contraction could be an optimal strategy for countering perturbation during reaching. In blocks of trials with different perturbation probabilities, they injected a torque pulse to the arm late in the reach. They found that subjects co-contracted their muscles in expectation of the perturbation, concluding that they used a feedforward strategy to overcome uncertainty. Our task could be viewed in a similar way where the sudden release of the handle is a random perturbation. Instead of simply countering the perturbation, our subjects co-activated their muscles pro-actively to optimize the entire movement. Our findings show that the feedforward strategy is employed more robustly as subjects learn the constraints of the task. Although beyond the scope of this study, reflexes associated with the rapid displacement of the arm may also be modulated predictively (Ludvig & Kearney, 2007) and act to restore stiffness disrupted by sudden changes in muscle length (Archambault et al., 2005; Crago et al., 1976; Liaw et al., 2008; Nichols & Houk, 1973, 1976; Reschechtko et al., 2019). However, since we observed stiffness-dependent co-activation before the release, a similar pattern of co-activation during the reflex, and because the impedance model was accurate through the entire task, these reflex contributions do not alter our main finding that subjects can accurately preset their stiffness to predict the mechanics of an upcoming movement.

## Conclusion

This work demonstrates that humans precisely and predictively tune their arm stiffness to accomplish a particularly challenging ballistic-release task. Our results show that subjects employed this strategy using the minimum possible stiffness and that the same strategy was used in different movement directions.

## Footnotes

1 The term “impedance” was introduced by Oliver Heaviside for linear electrical systems. It may be generalized rigorously to mechanical

2 Many of the developments in the algorithms for estimating impedance have been reported in a slightly different area of research, the study of the ankle joint (Guarín & Kearney, 2017; Lee et al., 2016; Ludvig et al., 2017; Rouse et al., 2014).

## References

Archambault, P. S., Mihaltchev, P., Levin, M. F., & Feldman, A. G. (2005). Basic elements of arm postural control analyzed by unloading. Experimental Brain Research, 164(2), 225–241. 10.1007/S00221-005-2245-6/FIGURES/11

Bennett, D. J., Hollerbach, J. M., Xu, Y., & Hunter, I. W. (1992). Time-varying stiffness of human elbow joint during cyclic voluntary movement. Experimental Brain Research, 88(2), 433–442. 10.1007/BF02259118/METRICS

Berret, B., Verdel, D., Burdet, E., & Jean, F. (2024). Co-contraction embodies uncertainty: An optimal feedforward strategy for robust motor control. PLOS Computational Biology, 20(11), e1012598. 10.1371/JOURNAL.PCBI.1012598

Burdet, E., Osu, R., Franklin, D. W., Milner, T. E., & Kawato, M. (2001). The central nervous system stabilizes unstable dynamics by learning optimal impedance. Nature 2001 414:6862, 414(6862), 446–449. 10.1038/35106566

Cannon, S. C., & Zahalak, G. I. (1982). The mechanical behavior of active human skeletal muscle in small oscillations. Journal of Biomechanics, 15(2), 111–121. 10.1016/0021-9290(82)90043-4

Crago, P. E., Houk, J. C., & Hasan, Z. (1976). Regulatory actions of human stretch reflex. 10.1152/Jn.1976.39.5.925, 39(5), 925–935. 10.1152/JN.1976.39.5.925

Feldman, A. G. (1966). On functional tuning of nervous system during controlled or preservation of stationary pose. 3. Mechanographic analysis of human performance of simple movement tasks. Biofizika, 11(4), 667–675. https://pubmed.ncbi.nlm.nih.gov/6000626/

Feldman, A. G. (1986). Once More on the Equilibrium-Point Hypothesis (λ Model) for Motor Control. Journal of Motor Behavior, 18(1), 17–54. 10.1080/00222895.1986.10735369

Franklin, D. W., Burdet, E., Osu, R., Kawato, M., & Milner, T. E. (2003a). Functional significance of stiffness in adaptation of multijoint arm movements to stable and unstable dynamics. Experimental Brain Research, 151(2), 145–157. 10.1007/S00221-003-1443-3/FIGURES/9

Franklin, D. W., Burdet, E., Osu, R., Kawato, M., & Milner, T. E. (2003b). Functional significance of stiffness in adaptation of multijoint arm movements to stable and unstable dynamics. Experimental Brain Research, 151(2), 145–157. 10.1007/S00221-003-1443-3/FIGURES/9

Franklin, D. W., Liaw, G., Milner, T. E., Osu, R., Burdet, E., & Kawato, M. (2007). Endpoint Stiffness of the Arm Is Directionally Tuned to Instability in the Environment. Journal of Neuroscience, 27(29), 7705–7716. 10.1523/JNEUROSCI.0968-07.2007

Garnier, H., Mensler, M., Richard, A., Garnier{, H., Mensler{, M., & Richard{, A. (2010). Continuous-time model identification from sampled data: Implementation issues and performance evaluation. 10.1080/0020717031000149636, 76(13), 1337–1357. 10.1080/0020717031000149636

Gomi, H., & Kawato, M. (1997). Human arm stiffness and equilibrium-point trajectory during multi-joint movement. Biological Cybernetics, 76(3), 163–171. 10.1007/S004220050329/METRICS

Graziano, M. S. A., & Gross, C. G. (1998). Spatial maps for the control of movement. Current Opinion in Neurobiology, 8(2), 195–201. 10.1016/S0959-4388(98)80140-2

Guarín, D. L., & Kearney, R. E. (2017). Estimation of time-varying, intrinsic and reflex dynamic joint stiffness during movement. Application to the ankle joint. Frontiers in Computational Neuroscience, 11, 257778. 10.3389/FNCOM.2017.00051/BIBTEX

Hinder, M. R., & Milner, T. E. (2003). The Case for an Internal Dynamics Model versus Equilibrium Point Control in Human Movement. The Journal of Physiology, 549(3), 953–963. 10.1113/JPHYSIOL.2002.033845

Hogan, N. (1980). Mechanical Impedance Control in Assistive Devices and Manipulators. In IEEE (Ed.), Proceedings of the IEEE Joint Automatic Control Conference.

Hogan, N. (1984). IMPEDANCE CONTROL: AN APPROACH TO MANIPULATION. Proceedings of the American Control Conference, 1, 304–313. 10.23919/ACC.1984.4788393

Hogan, N. (1985a). Impedance Control: An Approach to Manipulation: Part II—Implementation. Journal of Dynamic Systems, Measurement, and Control, 107(1), 8–16. 10.1115/1.3140713

Hogan, N. (1985b). Impedance Control: An Approach to Manipulation. Parts I-III. ASME: Journal of Dynamic Systems Measurement and Control, 107, 1–24.

Hogan, N. (1985c). The mechanics of multi-joint posture and movement control. Biological Cybernetics, 52(5), 315–331. 10.1007/BF00355754/METRICS

Hogan, N., & Sternad, D. (2012). Dynamic primitives of motor behavior. Biological Cybernetics, 106(11–12), 727–739. 10.1007/S00422-012-0527-1/METRICS

Kennedy, S. D., & Schwartz, A. B. (2019a). Distributed processing of movement signaling. Proceedings of the National Academy of Sciences, 116(52), 26266–26273. 10.1073/PNAS.1902296116

Kennedy, S. D., & Schwartz, A. B. (2019b). Stiffness as a control factor for object manipulation. Journal of Neurophysiology, 122(2), 707–720. 10.1152/JN.00372.2018/ASSET/IMAGES/LARGE/Z9K0081951510008.JPEG

Krutky, M. A., Trumbower, R. D., & Perreault, E. J. (2013). Influence of environmental stability on the regulation of end-point impedance during the maintenance of arm posture. Journal of Neurophysiology, 109(4), 1045–1054. 10.1152/JN.00135.2012/ASSET/IMAGES/LARGE/Z9K0041318040005.JPEG

Lackner, J. R., & Dizio, P. (1994). Rapid adaptation to Coriolis force perturbations of arm trajectory. 10.1152/Jn.1994.72.1.299, 72(1), 299–313. 10.1152/JN.1994.72.1.299

Lacquaniti, F., Carrozzo, M., & Borghese, N. A. (1993). Time-varying mechanical behavior of multijointed arm in man. 10.1152/Jn.1993.69.5.1443, 69(5), 1443–1464. 10.1152/JN.1993.69.5.1443

Lee, H., Rouse, E. J., & Krebs, H. I. (2016). Summary of Human Ankle Mechanical Impedance during Walking. IEEE Journal of Translational Engineering in Health and Medicine, 4. 10.1109/JTEHM.2016.2601613

Lemerle, S., Catalano, M. G., Bicchi, A., & Grioli, G. (2021). A Configurable Architecture for Two Degree-of-Freedom Variable Stiffness Actuators to Match the Compliant Behavior of Human Joints. Frontiers in Robotics and AI, 8, 614145. 10.3389/FROBT.2021.614145/BIBTEX

Lewis, G. N., Perreault, E. J., & MacKinnon, C. D. (2005). The influence of perturbation duration and velocity on the long-latency response to stretch in the biceps muscle. Experimental Brain Research, 163(3), 361–369. 10.1007/S00221-004-2182-9/FIGURES/6

Liaw, G., Franklin, D. W., Burdet, E., Kadi-Allah, A., & Kawato, M. (2008). Reflex contributions to the directional tuning of arm stiffness. Lecture Notes in Computer Science (Including Subseries Lecture Notes in Artificial Intelligence and Lecture Notes in Bioinformatics), 4984 LNCS(PART 1), 913–922. 10.1007/978-3-540-69158-7_94/COVER

Ljung, L. (2009). Experiments with Identification of Continuous Time Models. IFAC Proceedings Volumes, 42(10), 1175–1180. 10.3182/20090706-3-FR-2004.00195

Ludvig, D., & Kearney, R. E. (2007). Real-time estimation of intrinsic and reflex stiffness. IEEE Transactions on Biomedical Engineering, 54(10), 1875–1884. 10.1109/TBME.2007.894737

Ludvig, D., Plocharski, M., Plocharski, P., & Perreault, E. J. (2017). Mechanisms contributing to reduced knee stiffness during movement. Experimental Brain Research, 235(10), 2959–2970. 10.1007/S00221-017-5032-2/FIGURES/8

Mussa-Ivaldi, F. A., Hogan, N., & Bizzi, E. (1985). Neural, mechanical, and geometric factors subserving arm posture in humans. Journal of Neuroscience, 5(10), 2732–2743. 10.1523/JNEUROSCI.05-10-02732.1985

Nichols, T. R., & Houk, J. C. (1973). Reflex Compensation for Variations in the Mechanical Properties of a Muscle. Science, 181(4095), 182–184. 10.1126/SCIENCE.181.4095.182

Nichols, T. R., & Houk, J. C. (1976). Improvement in linearity and regulation of stiffness that results from actions of stretch reflex. 10.1152/Jn.1976.39.1.119, 39(1), 119–142. 10.1152/JN.1976.39.1.119

Orban, G. A., Claeys, K., Nelissen, K., Smans, R., Sunaert, S., Todd, J. T., Wardak, C., Durand, J. B., & Vanduffel, W. (2006). Mapping the parietal cortex of human and non-human primates. Neuropsychologia, 44(13), 2647–2667. 10.1016/J.NEUROPSYCHOLOGIA.2005.11.001

Paul, R. P. (1987). Problems and research issues associated with the hybrid control of force and displacement. Jet Propulsion Lab., California Inst. of Tech., Proceedings of the Workshop on Space Telerobotics, Volume 3.

Perreault, E. J., Kirsch, R. F., & Crago, P. E. (2001). Effects of voluntary force generation on the elastic components of endpoint stiffness. Experimental Brain Research, 141(3), 312–323. 10.1007/S002210100880/METRICS

Polit, A., & Bizzi, E. (1978). Processes Controlling Arm Movements in Monkeys. Science, 201(4362), 1235–1237. 10.1126/SCIENCE.99813

Polit, A., & Bizzi, E. (1979). Characteristics of motor programs underlying arm movements in monkeys. 10.1152/Jn.1979.42.1.183, 42(1), 183–194. 10.1152/JN.1979.42.1.183

Rancourt, D., & Hogan, N. (2009). The biomechanics of force production. Advances in Experimental Medicine and Biology, 629, 645–661. 10.1007/978-0-387-77064-2_35/FIGURES/35_5_978-0-387-77064-2

Reschechtko, S., Johansson, A. S., & Andrew Pruszynski, J. (2019). Maintaining arm control during self-triggered and unpredictable unloading perturbations. European Journal of Neuroscience, 50(10), 3531–3543. 10.1111/EJN.14479

Rouse, E. J., Hargrove, L. J., Perreault, E. J., & Kuiken, T. A. (2014). Estimation of human ankle impedance during the stance phase of walking. IEEE Transactions on Neural Systems and Rehabilitation Engineering, 22(4), 870–878. 10.1109/TNSRE.2014.2307256

Rudolph, K. S., Axe, M. J., & Snyder-Mackler, L. (2000). Dynamic stability after ACL injury: Who can hop? Knee Surgery, Sports Traumatology, Arthroscopy, 8(5), 262–269. 10.1007/S001670000130/METRICS

Schaal, S. (2006). Dynamic Movement Primitives -A Framework for Motor Control in Humans and Humanoid Robotics. Adaptive Motion of Animals and Machines, 261–280. 10.1007/4-431-31381-8_23

Schaal, S., Mohajerian, P., & Ijspeert, A. (2007). Dynamics systems vs. optimal control — a unifying view. Progress in Brain Research, 165, 425–445. 10.1016/S0079-6123(06)65027-9

Stout, D., & Chaminade, T. (2007). The evolutionary neuroscience of tool making. Neuropsychologia, 45(5), 1091–1100. 10.1016/J.NEUROPSYCHOLOGIA.2006.09.014

Takagi, A., Gomi, H., Burdet, E., & Koike, Y. (2022). A model predictive control strategy to regulate movements and interactions. BioRxiv, 2022.08.24.505193. 10.1101/2022.08.24.505193

Tee, K. P., Burdet, E., Chew, C. M., & Milner, T. E. (2004). A model of force and impedance in human arm movements. Biological Cybernetics, 90(5), 368–375. 10.1007/S00422-004-0484-4/METRICS

Trumbower, R. D., Krutky, M. A., Yang, B. S., & Perreault, E. J. (2009). Use of Self-Selected Postures to Regulate Multi-Joint Stiffness During Unconstrained Tasks. PLOS ONE, 4(5), e5411. 10.1371/JOURNAL.PONE.0005411

Tsuji, T. (1997). Human Arm Impedance in Multi-Joint Movements. Advances in Psychology, 119(C), 357–381. 10.1016/S0166-4115(97)80013-1

van de Ruit, M., Mugge, W., & Schouten, A. C. (2022). Quantifying Joint Stiffness During Movement: A Quantitative Comparison of Time-Varying System Identification Methods. Biosystems and Biorobotics, 28, 513–518. 10.1007/978-3-030-70316-5_82/FIGURES/3

Velliste, M., Perel, S., Spalding, M. C., Whitford, A. S., & Schwartz, A. B. (2008). Cortical control of a prosthetic arm for self-feeding. Nature 2008 453:7198, 453(7198), 1098–1101. 10.1038/nature06996

Walker, A. (2009). The strength of great apes and the speed of humans. Current Anthropology, 50(2), 229–234. 10.1086/592023/ASSET/IMAGES/LARGE/FG2.JPEG

Whitney, D. E. (1977). Force Feedback Control of Manipulator Fine Motions. Journal of Dynamic Systems, Measurement, and Control, 99(2), 91–97. 10.1115/1.3427095

Winter, D. A. (2009). Biomechanics and motor control of human movement. Wiley.

Young, P., Jakeman, A., Youngts, P., & Jakemant, A. (2007). Refined instrumental variable methods of recursive time-series analysis Part III. Extensions. 10.1080/00207178008961080, 31(4), 741–764. 10.1080/00207178008961080

