## Supplementary Material for "Tuning of Task-Relevant Stiffness in Multiple Directions"

[1]Dept. of Bioengineering, [2]Dept. of Neurobiology, Univ. of Pittsburgh, Pittsburgh, PA; [3]Dept. of Mechanical Eng., [4]Dept. of Brain and Cognitive Sci., MIT, Cambridge, MA

### Supplementary Material

*General subject information*

| SUBJECT # | GENDER | HEIGHT(CM) | WEIGHT(KG) |
|---|---|---|---|
| **1** | M | 170.2 | 63.2 |
| **2** | M | 175.0 | 65.7 |
| **3** | M | 182.9 | 85.8 |
| **4** | M | 160.0 | 50.2 |
| **5** | M | 183.0 | 82.0 |
| **6** | M | 172.0 | 77.8 |
| **7** | M | 180.0 | 90.0 |
| **8** | M | 183.0 | 78.5 |
| **9** | M | 182.0 | 81.3 |
| **10** | F | 163.0 | 60.0 |
| **11** | M | 180.0 | 75.0 |
| **12** | M | 182.0 | 113.0 |
| **13** | M | 188.0 | 101.5 |
| **14** | F | 163.0 | 55.2 |
| **15** | M | 180.3 | 65.0 |
| **16** | F | 162.6 | 62.1 |
| **17** | F | 160.0 | 58.0 |
| **18** | F | 157.5 | 58.5 |
| **19** | F | 168.0 | 57.3 |
| **20** | F | 162 | 46.4 |

*Success rate across subjects*

Success rate was defined as the number of successful trials (each subject needed to perform nine successful trials to complete each block) divided by the total number of trials in which the subject managed to complete the force hold stage of the task. Varying success rates were observed across subjects (Supplementary Figure 1) with a mean across directions of 64.10% $\pm$19%. One-way repeated measure ANOVA detected a statistically significant effect of direction on success rate ($F_{3,76} = 4.12, p = 0.0092$). Post-hoc pairwise comparisons showed a statistically significant difference between the +X and -Y direction ($p = 0.008$), while no significant difference was present between other directions.

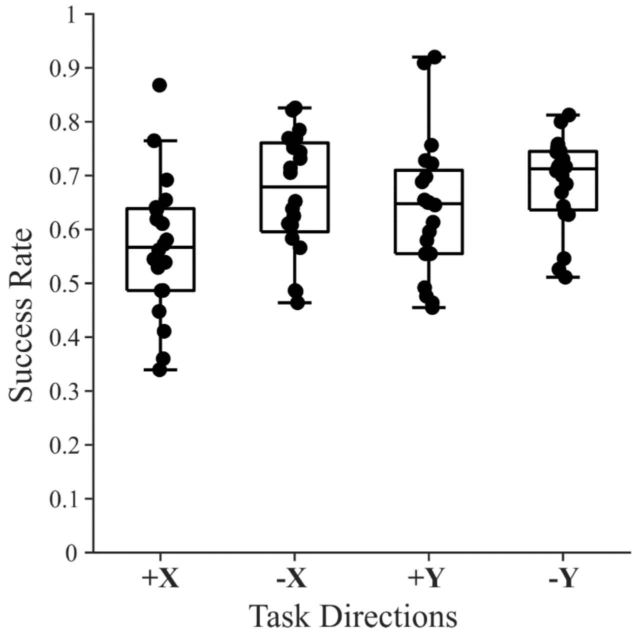


Supplementary Figure 1 – The success rate in different task directions. Dots show the task success rate of each subject. The box plot shows the median and the 25th and 75th percentile performance across subjects in each direction.

*2nd Order Model Fitting Performance*

Supplementary Table 1 reports the average fitting performance (FIT%) across trials and subjects in the 9 different conditions and 4 different directions.

| *FIT* % | $d = 2.5cm$ | $d = 5.0cm$ | $d = 7.5cm$ |
|---|---|---|---|
| $F = 15\ N$ | $92.3 \pm 2.4$ | $93.8 \pm 1.8$ | $95.5 \pm 1.0$ |
| $F = 20\ N$ | $90.4 \pm 5.4$ | $94.2 \pm 1.6$ | $94.9 \pm 1.4$ |
| $F = 25\ N$ | $88.1 \pm 6.4$ | $93.9 \pm 1.5$ | $95.3 \pm 1.0$ |

| *FIT* % | $d = 2.5cm$ | $d = 5.0cm$ | $d = 7.5cm$ |
|---|---|---|---|
| $F = 15\ N$ | $94.1 \pm 2.1$ | $95.9 \pm 1.4$ | $95.8 \pm 1.6$ |
| $F = 20\ N$ | $93.5 \pm 2.9$ | $95.7 \pm 1.5$ | $96.6 \pm 1.0$ |
| $F = 25\ N$ | $90.5 \pm 5.7$ | $96.4 \pm 1.0$ | $96.6 \pm 1.1$ |

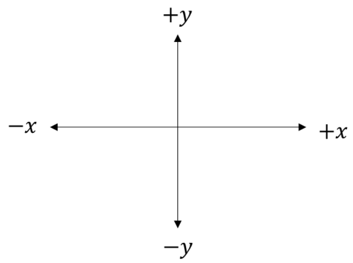


| *FIT* % | $d = 2.5cm$ | $d = 5.0cm$ | $d = 7.5cm$ |
|---|---|---|---|
| $F = 15\ N$ | $94.2 \pm 2.0$ | $94.5 \pm 2.0$ | $95.6 \pm 1.6$ |
| $F = 20\ N$ | $91.8 \pm 3.8$ | $94.4 \pm 1.9$ | $95.2 \pm 1.9$ |
| $F = 25\ N$ | $89.3 \pm 5.3$ | $94.3 \pm 1.5$ | $94.0 \pm 1.9$ |

| *FIT* % | $d = 2.5cm$ | $d = 5.0cm$ | $d = 7.5cm$ |
|---|---|---|---|
| $F = 15\ N$ | $93.3 \pm 1.7$ | $95.2 \pm 1.0$ | $95.1 \pm 1.1$ |
| $F = 20\ N$ | $92.4 \pm 3.2$ | $94.9 \pm 1.4$ | $95.1 \pm 0.9$ |
| $F = 25\ N$ | $89.5 \pm 5.5$ | $94.8 \pm 1.4$ | $96.0 \pm 0.9$ |

Supplementary Table 1 – The four different tables present the fitting performance across trials and subjects in the 9 force-displacement tested conditions for each movement direction. The reported numbers represent the mean and standard deviation of the fitting performance FIT%.

Supplementary Figure 2 shows the sensitivity of the FIT% to changes of $\pm$ 30% of the identified impedance parameters ($k_{est}, \beta_{est}, m_{est}$). In order to simplify the interpretation of sensitivity analysis, the estimated impedance values – as well as the FIT% – were normalized for each subject, direction, and condition.

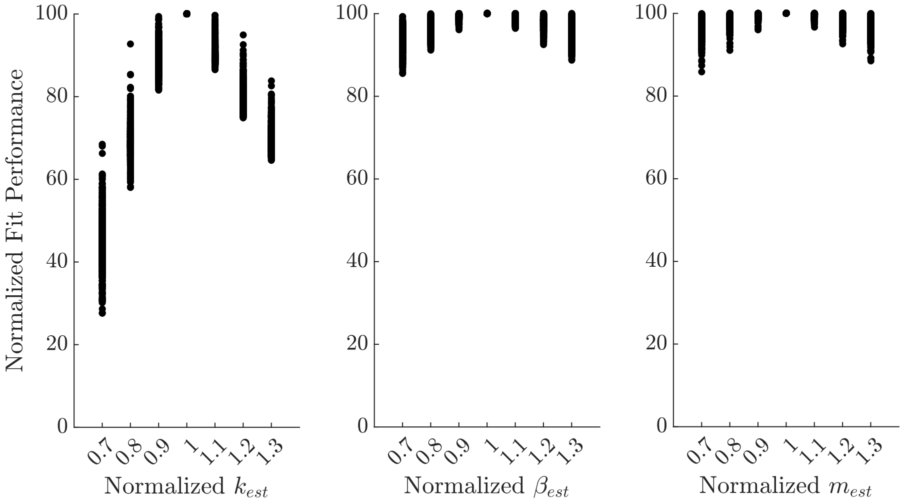


Supplementary Figure 2 – Sensitivity analysis of the normalized fitting performance (FIT%) to $\pm 30\%$ variations of the estimated impedance parameters ($k_{est}, \beta_{est}, m_{est}$) obtained from the 2nd order model system identification.

Supplementary Figure 3 reports the fitting performance differences between successful and failed trials as well as the difference in the stiffness estimates across all subjects and directions. The results confirm that 2nd order model was able to competently describe the arm end-effector behavior in all cases.

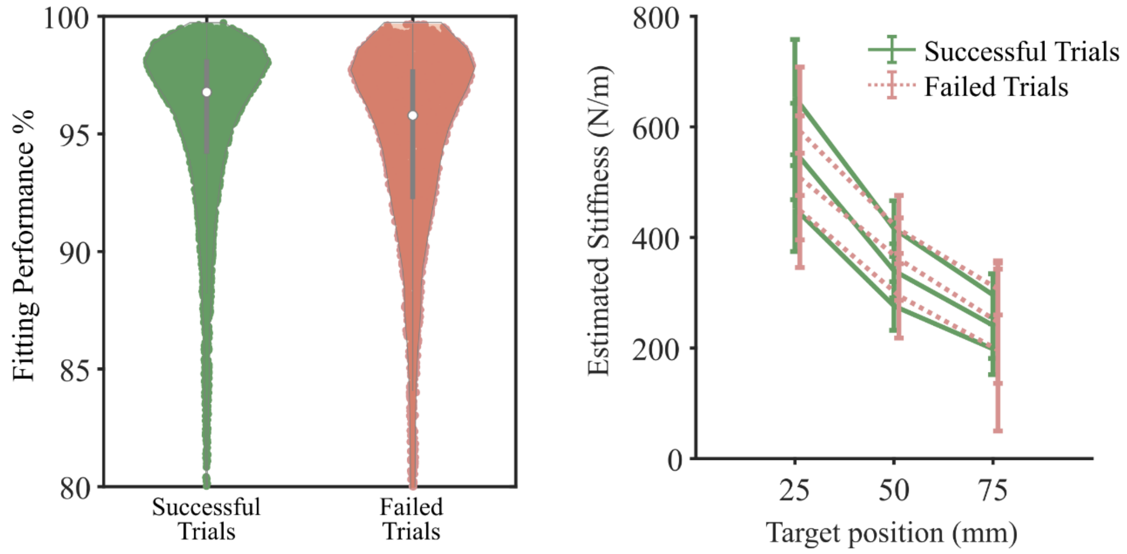

Supplementary Figure 3 – Fitting performance for the 2nd order model was found to be high for data from all trials irrespective of whether the subjects succeeded or failed. The estimated stiffness from the failed trials followed the same trend as the successful trials. This shows that the model could still fit the data. Trials were removed from the analysis not because the model did not fit but because subjects failed to comply with the task requirements.

Supplementary Figure 4 reports the estimated impedance parameters (mass, spring and damping) for a representative subject.

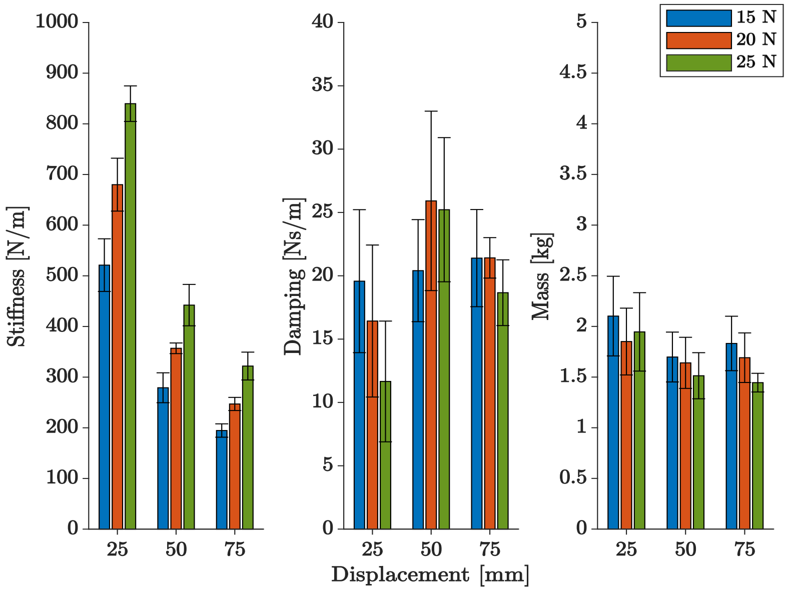


Supplementary Figure 4 – Estimated impedance parameters for a representative subject. For each panel, the x-axis shows the 3 different target displacements, while the 3 different colors represent the different target forces. Error bars indicate the standard deviation about the mean. The stiffness estimates (left panel) increased as force increased and decreased as the target displacement increased. In the middle panel, the damping coefficient appeared not to show any particular trend with respect to the different conditions. In the right panel, the mass was largely constant across different conditions.

Supplementary Figure 5 shows the estimated stiffness for all trials and all subjects for directions -X and -Y. The results, similar to Figure 8, show how subjects generated a task stiffness lower than the nominal task stiffness (solid blue bar) and close to the minimum allowed value required by the task.

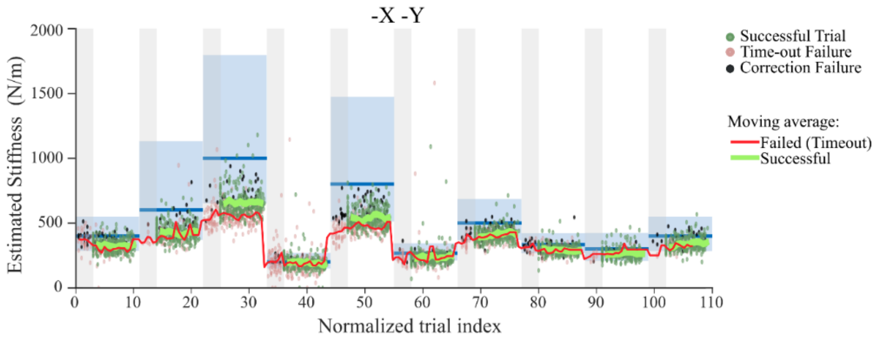


Supplementary Figure 5 – Estimated stiffness for all trials across sessions for directions -X and -Y (corresponding to +X, +Y directions shown in Figure 8). The figure follows the same format as Figure 8. The conditions were presented in the same order for these directions. White sections show the trials between the first and last successful trials in a condition. Gray sections show the failed trials between the last successful trial of one condition and the first successful trial of the next condition. Dots represent the estimated stiffness for each trial where green represents a successful trial, red represents failure due to time-out and black dots show failed trials due to corrections during movement. The blue bars show the range of nominal stiffness, $k_n = \frac{F_{tg}}{d_{tg}}$ and the blue rectangle shows the range of stiffness based on the allowed force and displacement thresholds. The red line shows the average estimated stiffness for failed (time-out) trials whereas the green solid lines show the average estimated stiffness for successful trials.

*Maximum Voluntary force*

We collected subjects' maximum voluntary force in different movement directions, calculated as the average force for 10s during which the subject was instructed to exert maximum isometric force. Maximum voluntary force varied from 46.1N to 211N, with forces in the left and back directions generally higher than in the front and right directions (Supplementary Figure 6A). All subjects' maximum voluntary force was greater than the highest force condition (25N) in the task. Maximum voluntary force was weakly but positively correlated (Supplementary Figure 6B) with body mass (R=0.5187).

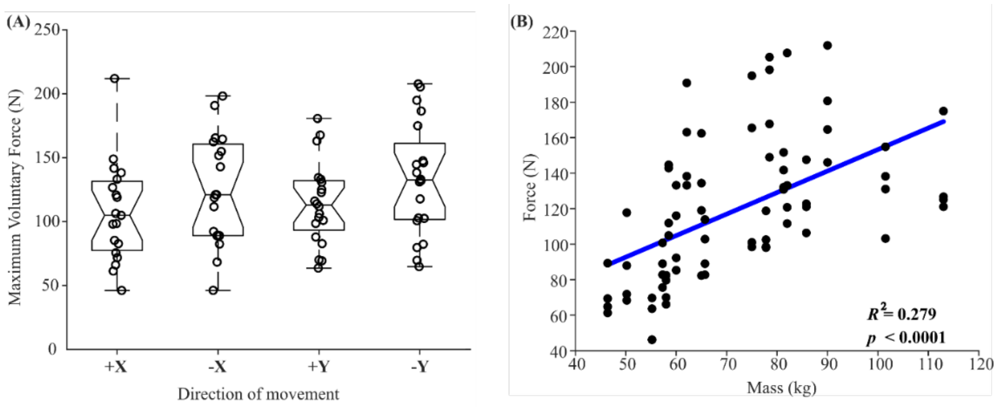

Supplementary Figure 6 – Maximum voluntary force. **A.** Maximum voluntary force across directions. Left and back direction forces were slightly higher than right and front. This might be because subjects had stronger pectoralis and biceps, which primarily exerted left and back forces, compared to posterior deltoid and triceps, which primarily generated right and front forces. **B.** Maximum voluntary force varied with subject mass. Subjects with greater mass tended to exert higher maximum voluntary force. The plot also reports the linear $R^2$ regression coefficient and a p-value, which indicated a slope significantly different from zero.

*Stiffness, Damping and Mass Estimates*

A two-way repeated measures ANOVA was performed to analyze the effect of force and displacement on stiffness, with trials as repeated measures and direction as a random factor. The analysis revealed that there was a statistically significant interaction between the effects of force and displacement ($F_{4,675} = 226.77, p < 0.001$). This interaction is evident in the top four plots of Figure 4, by the slight change in slope between the three target positions. Simple main-effects analysis showed that displacement had a statistically significant effect on stiffness in all pair-wise comparisons ($p < 0.05$). The same analysis applied to force showed that it also had a statistically significant effect on stiffness in all pair-wise comparisons ($p < 0.05$). These two effects are substantial and visibly evident in Figure 4 by the clear separation between the force and direction conditions. Supplementary Table 2 reports the complete post-hoc pairwise comparisons.

Nine two-way repeated-measures ANOVAs were performed to analyze the effect of direction on stiffness in the different force-displacement conditions. Trial was the repeated measure. The analysis revealed that there was a statistically significant effect ($p < 0.05$) of direction in all force-displacement conditions. Supplementary Table 3 reports the details of ANOVA results and all post-hoc pairwise comparisons. While significant, these differences were not substantial; the differences in estimated stiffness with respect to direction were contained within the range of expected target stiffness (light blue area in Figure 7) and accounted – on average – for only $8\% \pm 4\%$ of the minimum expected stiffness.

The top plots in Supplementary Figure 7 report the damping factor estimates across subjects in four different directions, while the bottom plots report the damping estimates in the nine different conditions with respect to movement direction.

Four two-way repeated measure ANOVAs were performed to analyze the effect of force and displacement on damping in the different directions. Trial was the repeated measure. The analysis revealed that there was a statistically significant interaction between the effects of force and displacement in all directions: +X($F_{4,162} = 37.57, p < 0.001$), -X ($F_{4,162} = 24.34, p < 0.001$), +Y ($F_{4,162} = 50.10, p = 0.015$), -Y ($F_{4,162} = 19.47, p < 0.001$). Simple main effects analysis showed that displacement had a statistically significant ($p < 0.05$) effect on damping in 81% of the pair-wise comparisons. Simple main effects analysis showed that force had a statistically significant ($p < 0.05$) effect on damping in 69% of the pair-wise comparisons. The Supplementary Material, Section E, reports the complete post-hoc pairwise comparisons in Supplementary Tables 4,5,6, and 7.

Nine two-way repeated-measure ANOVAs were performed to analyze the effect of direction on damping in the different force-displacement conditions. Trial was the repeated measure. The analysis revealed that there was a statistically significant effect ($p < 0.05$) of direction in all force-displacement conditions. Post-hoc pairwise comparisons showed that direction had a statistically significant ($p < 0.05$) effect on damping estimates in 85% of the pair-wise comparisons. The

Supplementary Material, Section E, reports the detailed ANOVA results and complete post-hoc pairwise comparisons in Supplementary Table 8.

The results show a high variability in the damping factor estimates, which appear noisy and without particular trend with respect to any of the analyzed parameters: direction of motion, target force, and target displacement. The counterintuitive significance can be explained by computing a linear fit of the damping factor estimates with respect to force and displacement. The 95% confidence intervals of the damping factor slope (with respect to both force and displacement) included zero in all but one case (23/24 slopes). This confirmed a high variability and insubstantial change of damping factor with respect to task conditions.

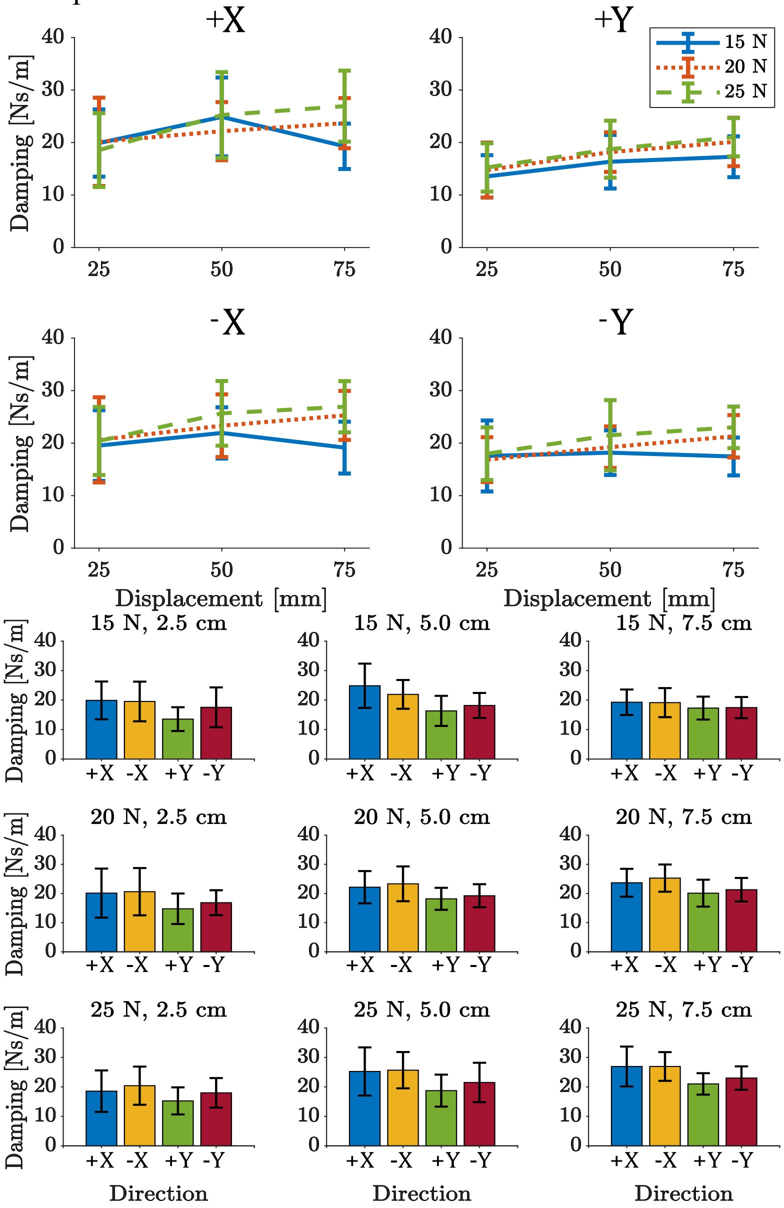


Supplementary Figure 7 – Arm damping variation across subjects. The top group of plots (4 plots) shows the damping coefficient change with respect to force and displacement in the four directions: +X (top left), +Y (top right), -X (bottom left), -Y (bottom right). For each panel, the 9 different force-displacement conditions are presented: the x-axis shows the different target displacements, 3 colored lines represent the different target forces. The bottom group of plots (9 plots) shows the damping change in the different force-displacement conditions with respect to the 4 directions of motion. The error bars in both plots show one standard deviation. (Complementary to Figure 7 in the main text with the same format).

To average the mass parameters across subjects, each estimated mass was normalized with respect to the subject's body weight. The estimated end-effector mass is a percentage of overall body weight. The normalized mass estimates in the four directions are shown in Supplementary Figure 8.

Four two-way repeated measure ANOVAs were performed to analyze the effect of force and displacement on mass in different directions. Trial was the repeated measure. The analysis revealed that there was a statistically significant interaction between the effects of force and displacement in all directions: +X($F_{4,162} = 18.72, p < 0.001$), -X ($F_{4,162} = 22.12, p < 0.001$), +Y ($F_{4,162} = 14.24, p < 0.001$), -Y ($F_{4,162} = 16.77, p < 0.001$). Simple main effects analysis showed that displacement had a statistically significant ($p < 0.05$) effect on mass in 81% of the pair-wise comparisons. Similarly, force had a statistically significant ($p < 0.05$) effect on mass in 47% of the pair-wise comparisons. The Supplementary Material, Section E, reports the complete post-hoc pairwise comparisons in Supplementary Tables 9,10,11, and 12.

Nine two-way repeated measure ANOVAs were performed to analyze the effect of direction on mass in the different force-displacement conditions. Trial was the repeated measure. The analysis revealed that there was a statistically significant effect ($p < 0.05$) of direction in all force-displacement conditions. Post-hoc pairwise comparisons showed that direction showed a statistically significant ($p < 0.05$) difference in mass estimates in 96% of the pair-wise comparisons. The Supplementary Material, Section E, reports the ANOVA detailed results and complete post-hoc pairwise comparisons in Supplementary Table 13.

Similar to the damping factor analysis, a linear fit of the mass estimates with respect to force and displacement showed a rate of change (slope) 95% Confidence Interval that included zero slope in all but three cases (21/24 slopes). This indicated that the mass estimates were statistically invariant with respect to target displacement and target force. A significant difference with respect to motion direction was evident. Specifically, the mass in the lateral direction ($\pm X$) was significantly higher than in the ($\pm$Y) frontal direction. This difference is attributable to the orientation of the hand mass ellipsoid, which is due to the arm inertia (refer to the Supplementary Material, Section G).

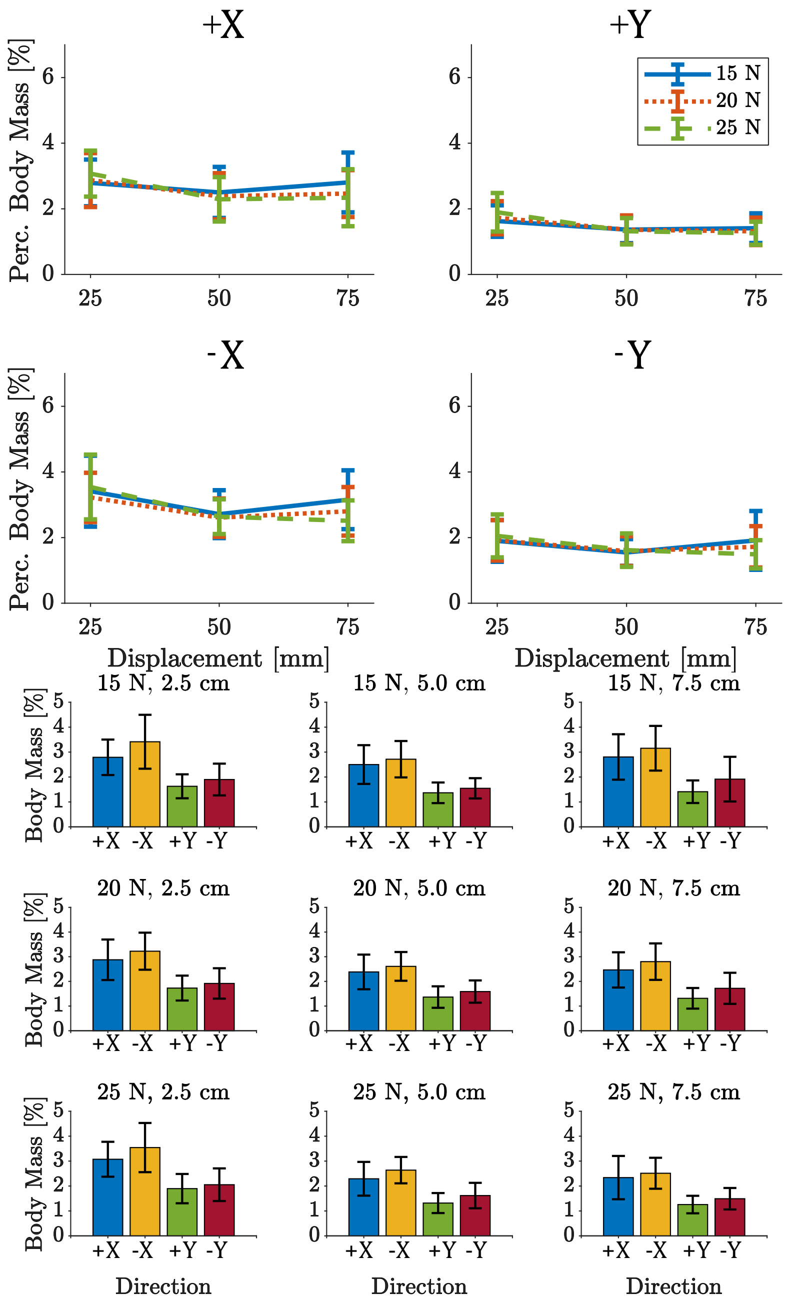


Supplementary Figure 8 – Arm end-effector mass variation across subjects. The top group of plots (4 plots) shows the mass coefficient (normalized by subject body weight) change with respect to force and displacement in the four directions: +X (top left), +Y (top right), -X (bottom left), -Y (bottom right). For each panel the 9 different force-displacement conditions are presented: the x-axis shows the different target displacements, 3 colored lines represent the different target forces. The bottom group of plots (9 plots) shows the mass change in the different force-displacement conditions with respect to the 4 directions of motion. The error bars in both plots show one standard deviation. (Complementary to Figure 7 in the main text with the same format).

*Two Degree of Freedom Arm Model*

A two DoF arm model was developed to analyze the effect of configuration changes on the end-effector mass and stiffness ellipses. The model, presented in Supplementary Figure 9, consisted of two rigid links representing the upper-arm and the forearm connected by means of simple rotational joints representing respectively the shoulder and elbow joints. The model considers the inertial properties of the human arm extracted from Winter (Winter, 2009). It did not include the gravitational load since the experiments were carried out in a plane perpendicular to the gravity vector and the arm was supported by a sling.

The equations of motions of such a system can be obtained by applying the Lagrange Equations, thus obtaining what follows:

$$M(\boldsymbol{\theta})\ddot{\boldsymbol{\theta}} + C(\boldsymbol{\theta},\dot{\boldsymbol{\theta}}) = \mathcal{F}$$

Where:

$$M(\boldsymbol{\theta}) = \begin{bmatrix} m_1L_{1G}^2 + m_2(L_1^2 + 2L_1L_{2G}\cos(\theta_2) + L_{2G}^2) + I_1 & m_2(L_1L_{2G}\cos(\theta_2) + L_{2G}^2) \\ m_2(L_1L_{2G}\cos(\theta_2) + L_{2G}^2) & m_2L_{2G}^2 + I_2 \end{bmatrix}$$

$$C(\boldsymbol{\theta},\dot{\boldsymbol{\theta}}) = \begin{bmatrix} -m_2L_1L_{2G}\left(2\dot{\theta}_1\dot{\theta}_2 + \dot{\theta}_2^2\right)\sin(\theta_2) \\ m_2L_1L_{2G}\dot{\theta}_1^2\sin(\theta_2) \end{bmatrix}$$

$$\mathcal{F} = \begin{bmatrix} \dfrac{\partial W}{\partial q_1} \\ \dfrac{\partial W}{\partial q_2} \end{bmatrix}$$

The geometrical and inertial quantities are summarized in the following list:

- Geometrical Quantities
  - $L_1$ → Shoulder-Elbow distance (experimentally measured)
  - $L_2$ → Elbow-Hand distance (experimentally measured)
  - $L_{1G} = 0.436L_1$ → Upper-arm Center of Mass (CoM) position (Winter, 2009)
  - $L_{2G} = 0.682L_2$ → Forearm Center of Mass (CoM) position (Winter, 2009)
- Inertial Quantities (Winter, 2009)
  - $m_1 = 0.028M$ → Upper-arm Mass with respect to Body Mass $M$
  - $m_2 = 0.022M$ → Forearm Mass with respect to Body Mass $M$
  - $I_1 = m_1\rho_{1G}^2$ → Upper-arm Moment of Inertia with respect to CoM
  - $I_2 = m_2\rho_{2G}^2$ → Forearm Moment of Inertia with respect to CoM
  - $\rho_{1G} = 0.322L_1$ → Upper-arm Radius of Gyration
  - $\rho_{2G} = 0.468L_2$ → Forearm Radius of Gyration

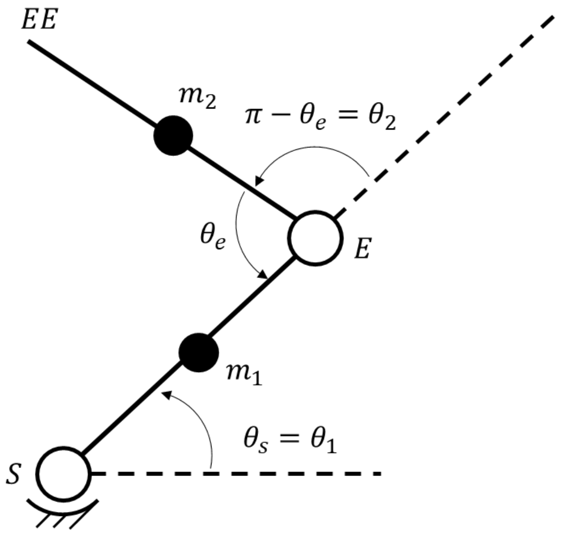


Supplementary Figure 9 – Schematic of the 2 DoF model of the arm. 'S' represents the shoulder joint, 'E' the elbow joint, 'EE' the position of the end-effector, '$\theta_s = \theta_1$' the shoulder angle, and '$\theta_e$' the elbow angle. $m_1$ is the mass of the upper arm, while $m_2$ is the mass of the forearm.

The mass matrix $M(\boldsymbol{\theta})$ represents the inertial properties of the model with respect to the configuration space i.e., the joint space. The same matrix can be expressed in terms of workspace coordinates, which in our case will be the end-effector planar coordinates (x,y), thus leading to the so-called end-effector mass matrix. From a mathematical standpoint, this is obtained as:

$$\Lambda(\boldsymbol{\theta}) = J^{-T}(\boldsymbol{\theta})M(\boldsymbol{\theta})J^{-1}(\boldsymbol{\theta})$$

Where $\Lambda(\boldsymbol{\theta})$ is the end-effector or spatial mass matrix and $J(\boldsymbol{\theta})$ is the Jacobian matrix computed as follows:

$$J(\boldsymbol{\theta}) = \begin{bmatrix} -L_1 \sin(\theta_1) - L_2 \sin(\theta_1 + \theta_2) & -L_2 \sin(\theta_1 + \theta_2) \\ -L_1 \cos(\theta_1) - L_2 \cos(\theta_1 + \theta_2) & -L_2 \cos(\theta_1 + \theta_2) \end{bmatrix}$$

In a similar way, the joint stiffness $R$ can be transformed to the end-effector stiffness matrix exploiting the Jacobian matrix. However, in this specific case, the joint stiffness matrix will also need to take into account also of the so-called kinematic stiffness $\Gamma = \frac{\delta J^T(\theta)}{\delta \theta} F$, which accounts for the apparent stiffness generated by external forces $F$(Tee et al., 2004).
Consequently, end-effector stiffness can be computed as:

$$K(\boldsymbol{\theta}) = J(\boldsymbol{\theta})^{-T} \cdot (R - \Gamma) \cdot J(\boldsymbol{\theta})^{-1}$$

*Stiffness tuning does not arise from bio-mechanical configuration*

To make sure that the observed stiffness tuning did not arise from the difference in the bio-mechanical configurations during movement in the different directions, we estimated the endpoint stiffness using the joint configuration data collected during the experiment. For stiffness coefficients, the model took a constant joint impedance and the joint position collected by the motion-tracking system (Optotrak 3020, Northern Digital, Canada) as input and gave endpoint impedance as output. For mass coefficients, the model took each subject's body-mass index (BMI) and joint positions as input and gave end-effector mass as output (each body segment mass was assumed according to Winter (Winter, 2009)).

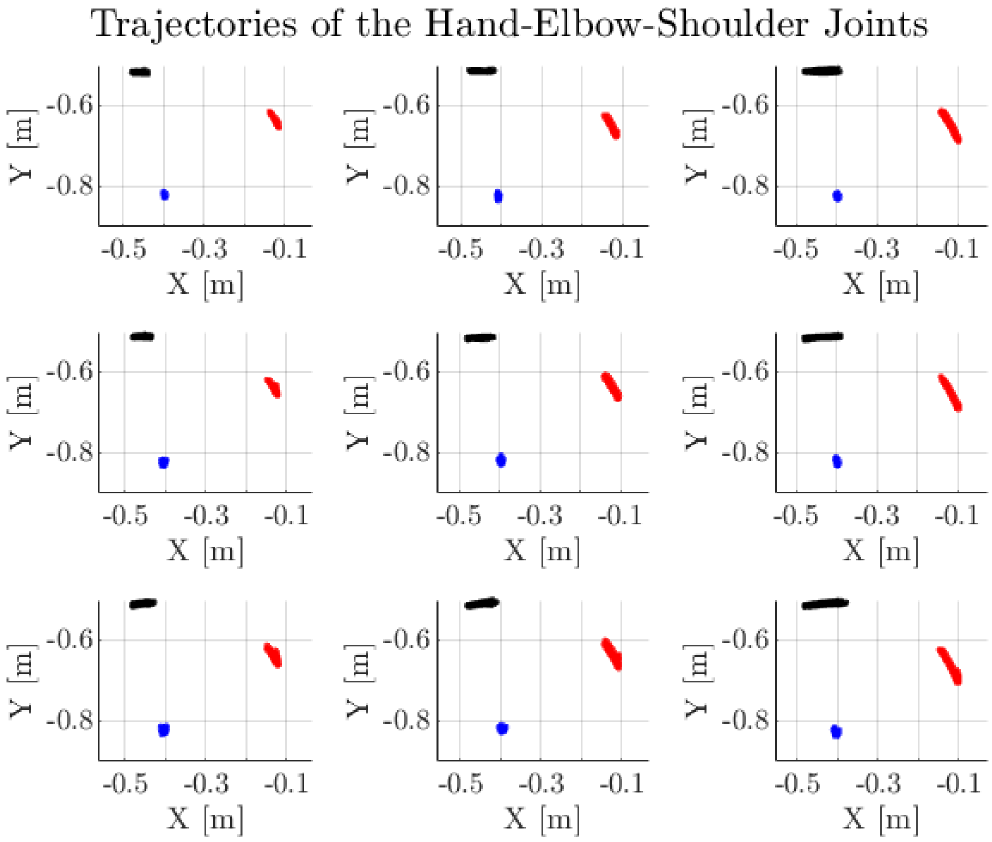


Supplementary Figure 10 – Trajectories of the hand (black dots), elbow (red dots), and shoulder (blue dots) markers in the horizontal (x,y) plane for a representative subject. The 9 panels represent the nine different force-displacement conditions performed during the ballistic release experiment. The target force increases from the first to the last row, while the target displacement increases from the first to the last column.

We calculated the predicted endpoint stiffness as a series of ellipses, depending on the different joint locations, and plotted the start and final locations with red and black separately. We found that the endpoint stiffness before release consisted of different force and position target conditions, while the endpoint stiffness at the end of the movement depended only on the target position. The fact that the kinematic stiffness did not change across different force levels with the same target indicated that the subjects tuned their joint stiffness according to different force levels. Furthermore, the kinematic stiffness failed to predict the endpoint stiffness variation with different targets, but the same force indicated that subjects tuned their joint stiffness corresponding to different target displacement levels.

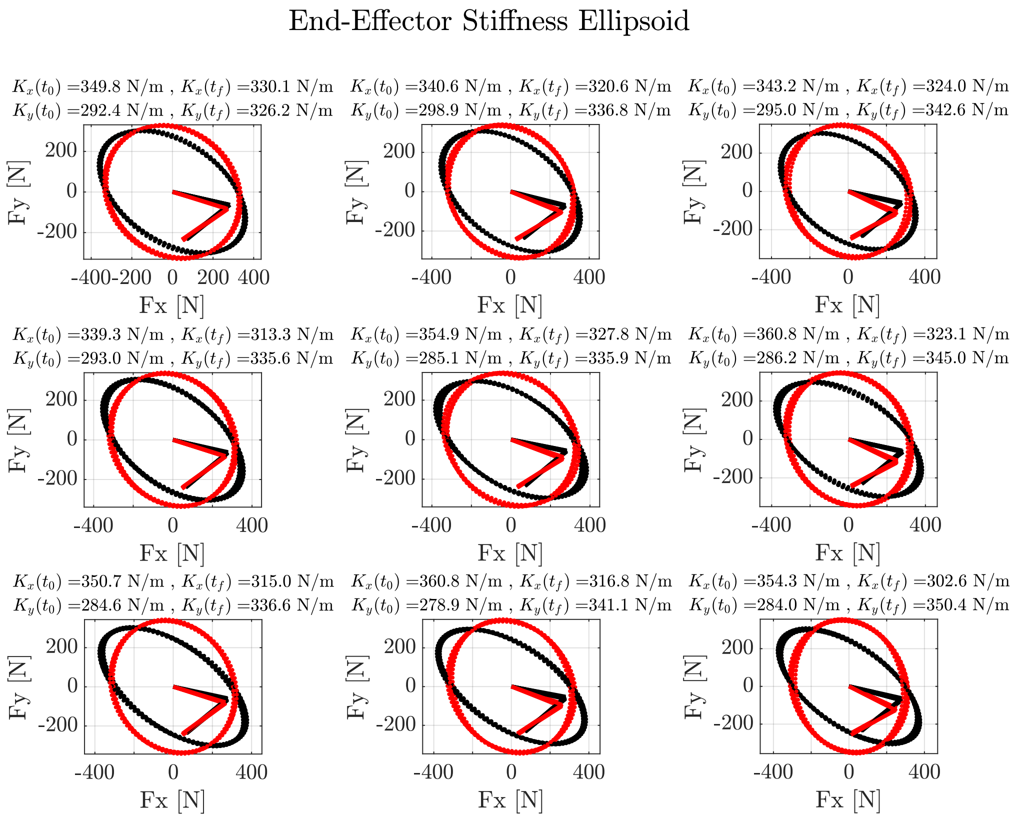


Supplementary Figure 11 – The kinematic stiffness prediction in different task conditions obtained using the two degree of freedom arm model. The 9 panels represent the 9 nine different force-displacement conditions performed during the ballistic release experiment. The target force increases from the first to the last row, while the target displacement increases from the first to the last column. For each panel two ellipses are presented: the black ellipse represents the end-effector stiffness ellipse during force hold, while the red ellipse shows the end-effector stiffness ellipse during position hold. The segments in each figure are added to show the difference in position of the arm model. On top of each figure the values of the resulting end-effector stiffness in both the X (lateral) and Y (frontal) directions, respectively $K_x$ and $K_y$, are reported both during the force hold phase ($t_0$) and in the final position hold phase ($t_f$).

The endpoint mass was estimated in the same way. Each body segment mass and center of mass was estimated from participants' height and weight. The endpoint mass was calculated as described above. The ellipses from different conditions only changed with different movement positions. The fact that the mass prediction from the kinematic model overlapped the mass coefficient estimated from the $2^{nd}$-order model validates the estimates from our $2^{nd}$-order model.

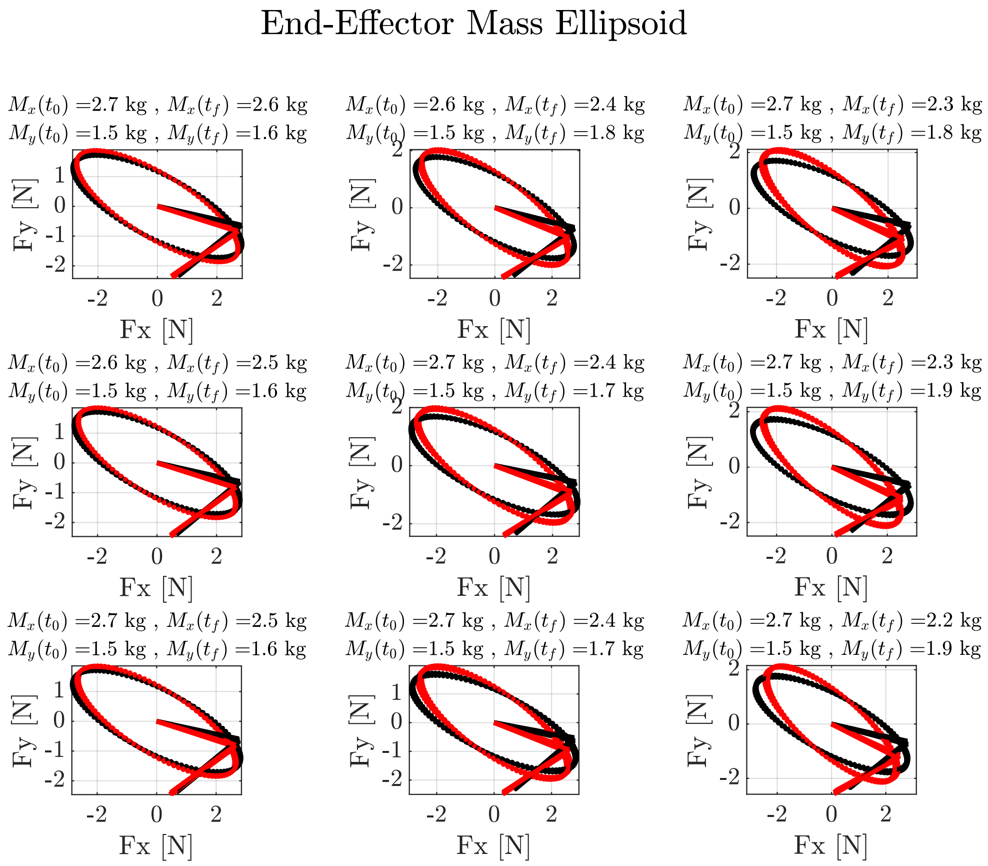


Supplementary Figure 12 – The endpoint mass predictions in different task conditions were obtained using the two degree of freedom arm model. The 9 panels represent the 9 nine different force-displacement conditions performed during the ballistic release experiment. The target force increases from the first to the last row, while the target displacement increases from the first to the last column. For each panel two ellipses are presented: the black ellipse represents the end-effector mass ellipse during force hold, while the red ellipse shows the end-effector mass ellipse during position hold. The segments in each figure are added to show the difference in position of the arm model. On top of each figure the values of the resulting end-effector mass in both the X (lateral) and Y (frontal) directions, respectively $M_x$ and $M_y$, are reported both during the force hold phase ($t_0$) and in the final position hold phase ($t_f$).

#### *EMG data analysis*

Supplementary Figure 13A shows the average normalized EMG across all subjects from the three antagonist muscle pairs during the force hold phase of the task for the highest stiffness condition in the four task directions. The wrist muscles were active in all directions whereas the Biceps-Triceps and the deltoids were active only in certain directions. Moreover, the CAI was found to be strongly positively correlated with stiffness when the muscles were active (Supplementary Figure 13B).

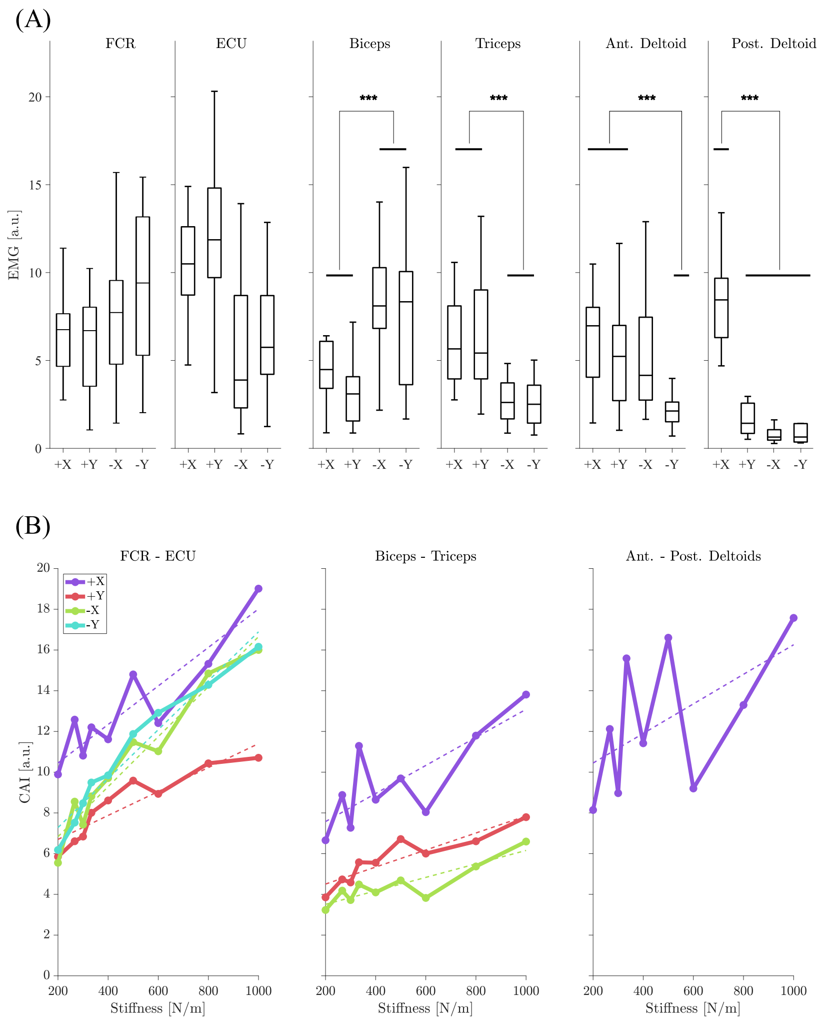


Supplementary Figure 13 - EMG dependency on movement direction. **A.** Average EMG across all subjects from the force-hold phase for all muscles for the four task directions. The FCR and ECU were active in all directions. Asterisk indicate statistical significance of the difference in activity across directions (***: $p < 0.001$). The biceps and triceps were active in opposite directions. The anterior deltoid was active in three directions, while the posterior deltoid was active only in one direction. **B.** CAI of different muscle pairs for the directions in which the muscles pairs were active. The CAI was strongly and positively correlated with stiffness in the active directions.

Recorded EMG from the terminal position hold phase of the task was used to compute the co-activation index (CAI), which was then regressed against the three task parameters (K, F, and X) across all subjects and directions. Similar to the pre-movement phase, the co-activation of wrist muscles was found to be strongly correlated with stiffness but only weakly with force or displacement.

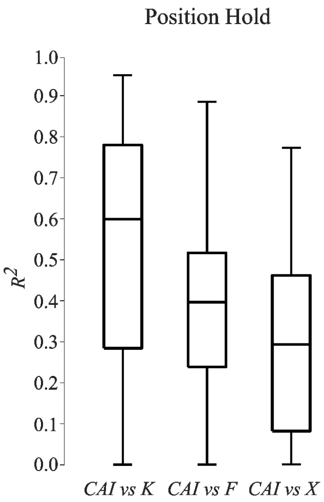


Supplementary Figure 14 – Co-activation index from average EMG activity recorded during the terminal position hold phase of the task was regressed against the various task parameters (similar to those shown in Figures 9D and Figure 10D). We found that the goodness of fit increased to the force-hold phase levels, especially for the stiffness parameter, suggesting that subjects maintained their co-activation of wrist muscles even when they reached the target location.

### *Post-hoc t-tests Complete Results*

The post-hoc pairwise comparisons were performed when significant interaction was observed in any of the performed repeated measure ANOVAs. The following is a complete report of the pairwise comparisons divided per dependent variable.

#### *Stiffness*

##### *Effect of Target Force and Target Displacement*

Supplementary Table 2 - Pairwise comparisons of stiffness estimates with respect to target force and target displacement. The table reports the Bonferroni adjusted p-values for multiple comparisons. The significance was set to p=0.05. The left part of the table reports the comparisons between target displacements at the different force levels. The right part of the table reports the comparison between target forces at different displacement levels. The asterisk * denotes a significant difference.

| | **d=2.5cm/5.0cm** | **d=5.0/7.5cm** | **d=2.5cm/7.5cm** | | **F=15N/20N** | **F=20N/25N** | **F=15N/25N** |
|---|---|---|---|---|---|---|---|
| **F=15N** | $p<0.001$* | $p<0.001$* | $p<0.001$* | **d=2.5cm** | $p<0.001$* | $p<0.001$* | $p<0.001$* |
| **F=20N** | $p<0.001$* | $p<0.001$* | $p<0.001$* | **d=5.0cm** | $p<0.001$* | $p<0.001$* | $p<0.001$* |
| **F=25N** | $p<0.001$* | $p<0.001$* | $p<0.001$* | **d=7.5cm** | $p<0.001$* | $p<0.001$* | $p<0.001$* |

##### *Effect of Direction*

Supplementary Table 3 - This table reports the repeated measure ANOVA results (second column) and the post-hoc pairwise comparisons of stiffness estimates with respect to direction. The pair-wise comparisons were Bonferroni adjusted for multiple comparisons. The significance was set to p=0.05. The asterisk * denotes a significant difference.

| | **ANOVA** | **(+X, -X)** | **(+X, +Y)** | **(+X, -Y)** | **(-X, +Y)** | **(-X, -Y)** | **(+Y, -Y)** |
|---|---|---|---|---|---|---|---|
| **(15N,2.5cm)** | $F = 27.01$ $p < 0.001$* | $p<0.001$* | $p=1.000$ | $p<0.001$* | $p<0.001$* | $p=0.013$* | $p<0.001$* |
| **(15N,5.0cm)** | $F = 46.78$ | $p<0.001$* | $p=0.035$* | $p<0.001$* | $p<0.001$* | $p<0.001$* | $p<0.001$* |

| | | | | | | | |
|---|---|---|---|---|---|---|---|
| | $p < 0.001$* | | | | | | |
| **(15N,7.5cm)** | $F = 56.15$ $p < 0.001$* | $p$<0.001* | $p$=1.000 | $p$<0.001* | $p$<0.001* | $p$=1.000 | $p$<0.001* |
| **(20N,2.5cm)** | $F = 20.06$ $p < 0.001$* | $p$<0.001* | $p$<0.001* | $p$<0.001* | $p$=1.000 | $p$=1.000 | $p$=1.000 |
| **(20N,5.0cm)** | $F = 5.79$ $p < 0.001$* | $p$<0.001* | $p$=1.000 | $p$=1.000 | $p$=0.102 | $p$=0.005* | $p$=1.000 |
| **(20N,7.5cm)** | $F = 18.83$ $p < 0.001$* | $p$<0.001* | $p$=0.608 | $p$<0.001* | $p$<0.001* | $p$=0.414 | $p$=0.016* |
| **(25N,2.5cm)** | $F = 10.62$ $p < 0.001$* | $p$=0.015* | $p$<0.001* | $p$=0.042* | $p$=0.015* | $p$=1.000 | $p$=0.135 |
| **(25N,5.0cm)** | $F = 12.22$ $p < 0.001$* | $p$<0.001* | $p$=0.103 | $p$=0.854 | $p$<0.001* | $p$<0.001* | $p$=1.000 |
| **(25N,7.5cm)** | $F = 31.67$ $p < 0.001$* | $p$<0.001* | $p$=0.418 | $p$<0.001* | $p$<0.001* | $p$=1.000 | $p$<0.001* |

*Damping*

*Effect of Target Force and Target Displacement: +X*

Supplementary Table 4 - Pairwise comparisons of damping estimates in +X direction with respect to target force and target displacement. The table reports the Bonferroni adjusted p-values for multiple comparisons. The significance was set to p=0.05. The left part of the table reports the comparisons between target displacements at the different force levels. The right part of the table reports the comparison between target forces at different displacement levels. The asterisk * denotes a significant difference.

| | **d=2.5cm/5.0cm** | **d=5.0/7.5cm** | **d=2.5cm/7.5cm** | | **F=15N/20N** | **F=20N/25N** | **F=15N/25N** |
|---|---|---|---|---|---|---|---|
| **F=15N** | $p$<0.001* | $p$<0.001* | $p$=0.842 | **d=2.5cm** | $p$=1.000 | $p$=0.021* | $p$=0.110 |
| **F=20N** | $p$=0.011* | $p$=0.005* | $p$<0.001* | **d=5.0cm** | $p$<0.001* | $p$<0.001* | $p$=1.000 |
| **F=25N** | $p$<0.001* | $p$=0.052 | $p$<0.001* | **d=7.5cm** | $p$<0.001* | $p$<0.001* | $p$<0.001* |

*Effect of Target Force and Target Displacement: -X*

Supplementary Table 5 - Pairwise comparisons of damping estimates in -X direction with respect to target force and target displacement. The table reports the Bonferroni adjusted p-values for multiple comparisons. The significance was set to p=0.05. The left part of the table reports the comparisons between target displacements at the different force levels. The right part of the table reports the comparison between target forces at different displacement levels. The asterisk * denotes a significant difference.

| | **d=2.5cm/5.0cm** | **d=5.0/7.5cm** | **d=2.5cm/7.5cm** | | **F=15N/20N** | **F=20N/25N** | **F=15N/25N** |
|---|---|---|---|---|---|---|---|
| **F=15N** | $p$<0.001* | $p$<0.001* | $p$=1.000 | **d=2.5cm** | $p$=0.347 | $p$=1.000 | $p$=0.377 |
| **F=20N** | $p$<0.001* | $p$<0.001* | $p$<0.001* | **d=5.0cm** | $p$=0.005* | $p$<0.001* | $p$<0.001* |
| **F=25N** | $p$<0.001* | $p$=0.020* | $p$<0.001* | **d=7.5cm** | $p$<0.001* | $p$<0.001* | $p$=0.001* |

*Effect of Target Force and Target Displacement: +Y*

Supplementary Table 6 - Pairwise comparisons of damping estimates in +Y direction with respect to target force and target displacement. The table reports the Bonferroni adjusted p-values for multiple comparisons. The significance was set to p=0.05. The left part of the table reports the comparisons between target displacements at the different force levels. The right part of the table reports the comparison between target forces at different displacement levels. The asterisk * denotes a significant difference.

| | **d=2.5cm/5.0cm** | **d=5.0/7.5cm** | **d=2.5cm/7.5cm** | | **F=15N/20N** | **F=20N/25N** | **F=15N/25N** |
|---|---|---|---|---|---|---|---|
| **F=15N** | $p$<0.001* | $p$=0.188 | $p$<0.001* | **d=2.5cm** | $p$=0.020* | $p$=0.554 | $p$<0.001* |
| **F=20N** | $p$<0.001* | $p$<0.001* | $p$<0.001* | **d=5.0cm** | $p$<0.001* | $p$=0.581 | $p$<0.001* |
| **F=25N** | $p$<0.001* | $p$<0.001* | $p$<0.001* | **d=7.5cm** | $p$<0.001* | $p$=0.087 | $p$<0.001* |

*Effect of Target Force and Target Displacement: -Y*

Supplementary Table 7 - Pairwise comparisons of damping estimates in -Y direction with respect to target force and target displacement. The table reports the Bonferroni adjusted p-values for multiple comparisons. The significance was set to p=0.05. The left part of the table reports the comparisons between target displacements at the different force levels. The right part of the table reports the comparison between target forces at different displacement levels. The asterisk * denotes a significant difference.

| | d=2.5cm/5.0cm | d=5.0/7.5cm | d=2.5cm/7.5cm | | F=15N/20N | F=20N/25N | F=15N/25N |
|---|---|---|---|---|---|---|---|
| F=15N | p=0.714 | p=0.157 | p=1.000 | d=2.5cm | p=0.594 | p=0.025* | p=1.000 |
| F=20N | p<0.001* | p<0.001* | p<0.001* | d=5.0cm | p=0.042* | p<0.001* | p<0.001* |
| F=25N | p<0.001* | p=0.016* | p<0.001* | d=7.5cm | p<0.001* | p<0.001* | p<0.001* |

*Effect of Direction*

Supplementary Table 8 – The table reports the repeated measures ANOVA results (second column) and the post-hoc pairwise comparisons of damping estimates with respect to direction. The pair-wise comparisons were Bonferroni adjusted for multiple comparisons. The significance was set to p=0.05. The asterisk * denotes a significant difference.

| | ANOVA | (+X, -X) | (+X, +Y) | (+X, -Y) | (-X, +Y) | (-X, -Y) | (+Y, -Y) |
|---|---|---|---|---|---|---|---|
| (15N,2.5cm) | $F = 49.55$ $p < 0.001$* | p=1.000 | p<0.001* | p=0.004* | p<0.001* | p=0.007* | p<0.001* |
| (15N,5.0cm) | $F = 93.59$ $p < 0.001$* | p<0.001* | p<0.001* | p<0.001* | p<0.001* | p<0.001* | p=0.002* |
| (15N,7.5cm) | $F = 12.56$ $p < 0.001$* | p=1.000 | p<0.001* | p<0.001* | p=0.001* | p<0.001* | p=1.000 |
| (20N,2.5cm) | $F = 31.56$ $p < 0.001$* | p=1.000 | p<0.001* | p<0.001* | p<0.001* | p<0.001* | p<0.001* |
| (20N,5.0cm) | $F = 102.67$ $p < 0.001$* | p=0.304 | p<0.001* | p<0.001* | p<0.001* | p<0.001* | p<0.001* |
| (20N,7.5cm) | $F = 53.05$ $p < 0.001$* | p=0.003* | p<0.001* | p<0.001* | p<0.001* | p<0.001* | p=0.025* |
| (25N,2.5cm) | $F = 31.35$ $p < 0.001$* | p=0.006* | p<0.001* | p=1.000 | p<0.001* | p<0.001* | p<0.001* |
| (25N,5.0cm) | $F = 51.22$ $p < 0.001$* | p=1.000 | p<0.001* | p<0.001* | p<0.001* | p<0.001* | p<0.001* |
| (25N,7.5cm) | $F = 67.42$ $p < 0.001$* | p=1.000 | p<0.001* | p<0.001* | p<0.001* | p<0.001* | p<0.001* |

*Mass*

*Effect of Target Force and Target Displacement: +X*

Supplementary Table 9 - Pairwise comparisons of mass estimates in +X direction with respect to target force and target displacement. The table reports the Bonferroni adjusted p-values for multiple comparisons. The significance was set to p=0.05. The left part of the table reports the comparisons between target displacements at the different force levels. The right part of the table reports the comparison between target forces at different displacement levels. The asterisk * denotes a significant difference.

| | d=2.5cm/5.0cm | d=5.0/7.5cm | d=2.5cm/7.5cm | | F=15N/20N | F=20N/25N | F=15N/25N |
|---|---|---|---|---|---|---|---|
| F=15N | p<0.001* | p<0.001* | p=1.000 | d=2.5cm | p=0.595 | p=0.003* | p<0.001* |
| F=20N | p<0.001* | p=0.657 | p<0.001* | d=5.0cm | p=0.153 | p=0.277 | p=0.004* |
| F=25N | p<0.001* | p=1.000 | p<0.001* | d=7.5cm | p<0.001* | p=0.203 | p<0.001* |

*Effect of Target Force and Target Displacement: -X*

Supplementary Table 10 - Pairwise comparisons of mass estimates in -X direction with respect to target force and target displacement. The table reports the Bonferroni adjusted p-values for multiple comparisons. The significance was set to p=0.05. The left part of the table reports the comparisons between target displacements at the different force levels. The right part of the table reports the comparison between target forces at different displacement levels. The asterisk * denotes a significant difference.

| | d=2.5cm/5.0cm | d=5.0/7.5cm | d=2.5cm/7.5cm | | F=15N/20N | F=20N/25N | F=15N/25N |
|---|---|---|---|---|---|---|---|
| F=15N | p<0.001* | p<0.001* | p=0.013* | d=2.5cm | p=0.036* | p<0.001* | p=0.232 |
| F=20N | p<0.001* | p=0.013* | p<0.001* | d=5.0cm | p=0.165 | p=1.000 | p=0.496 |
| F=25N | p<0.001* | p=0.019* | p<0.001* | d=7.5cm | p<0.001* | p<0.001* | p<0.001* |

*Effect of Target Force and Target Displacement: +Y*

Supplementary Table 11 - Pairwise comparisons of mass estimates in +Y direction with respect to target force and target displacement. The table reports the Bonferroni adjusted p-values for multiple comparisons. The significance was set to p=0.05. The left part of the table reports the comparisons between target displacements at the different force levels. The right part of the table reports the comparison between target forces at different displacement levels. The asterisk * denotes a significant difference.

| | d=2.5cm/5.0cm | d=5.0/7.5cm | d=2.5cm/7.5cm | | F=15N/20N | F=20N/25N | F=15N/25N |
|---|---|---|---|---|---|---|---|
| **F=15N** | *p*<0.001* | *p*=0.724 | *p*<0.001* | **d=2.5cm** | *p*=0.099 | *p*<0.001* | *p*<0.001* |
| **F=20N** | *p*<0.001* | *p*=0.711 | *p*<0.001* | **d=5.0cm** | *p*=1.000 | *p*=0.562 | *p*=0.420 |
| **F=25N** | *p*<0.001* | *p*=0.265 | *p*<0.001* | **d=7.5cm** | *p*=0.079 | *p*=0.378 | *p*<0.001* |

*Effect of Target Force and Target Displacement: -Y*

Supplementary Table 12 - Pairwise comparisons of mass estimates in -Y direction with respect to target force and target displacement. The table reports the Bonferroni adjusted p-values for multiple comparisons. The significance was set to p=0.05. The left part of the table reports the comparisons between target displacements at the different force levels. The right part of the table reports the comparison between target forces at different displacement levels. The asterisk * denotes a significant difference.

| | d=2.5cm/5.0cm | d=5.0/7.5cm | d=2.5cm/7.5cm | | F=15N/20N | F=20N/25N | F=15N/25N |
|---|---|---|---|---|---|---|---|
| **F=15N** | *p*<0.001* | *p*<0.001* | *p*=1.000 | **d=2.5cm** | *p*=1.000 | *p*=0.008* | *p*=0.027* |
| **F=20N** | *p*<0.001* | *p*=0.035* | *p*<0.001* | **d=5.0cm** | *p*=0.875 | *p*=1.000 | *p*=0.321 |
| **F=25N** | *p*<0.001* | *p*=0.007* | *p*<0.001* | **d=7.5cm** | *p*=0.064 | *p*<0.001* | *p*<0.001* |

*Effect of Direction*

Supplementary Table 13 - The table reports the repeated measure ANOVA results (second column) and the post-hoc pairwise comparisons of mass estimates with respect to direction. The pair-wise comparisons were Bonferroni adjusted for multiple comparisons. The significance was set to p=0.05. The asterisk * denotes a significant difference.

| | ANOVA | (+X, -X) | (+X, +Y) | (+X, -Y) | (-X, +Y) | (-X, -Y) | (+Y, -Y) |
|---|---|---|---|---|---|---|---|
| **(15N,2.5cm)** | $F = 245.49$ $p < 0.001$* | *p*<0.001* | *p*<0.001* | *p*<0.001* | *p*<0.001* | *p*<0.001* | *p*<0.001* |
| **(15N,5.0cm)** | $F = 234.64$ $p < 0.001$* | *p*=0.017* | *p*<0.001* | *p*<0.001* | *p*<0.001* | *p*<0.001* | *p*<0.001* |
| **(15N,7.5cm)** | $F = 174.88$ $p < 0.001$* | *p*<0.001* | *p*<0.001* | *p*<0.001* | *p*<0.001* | *p*<0.001* | *p*<0.001* |
| **(20N,2.5cm)** | $F = 261.62$ $p < 0.001$* | *p*<0.001* | *p*<0.001* | *p*<0.001* | *p*<0.001* | *p*<0.001* | *p*=0.005* |
| **(20N,5.0cm)** | $F = 263.37$ $p < 0.001$* | *p*=0.001* | *p*<0.001* | *p*<0.001* | *p*<0.001* | *p*<0.001* | *p*<0.001* |
| **(20N,7.5cm)** | $F = 209.95$ $p < 0.001$* | *p*<0.001* | *p*<0.001* | *p*<0.001* | *p*<0.001* | *p*<0.001* | *p*<0.001* |
| **(25N,2.5cm)** | $F = 228.07$ $p < 0.001$* | *p*<0.001* | *p*<0.001* | *p*<0.001* | *p*<0.001* | *p*<0.001* | *p*=0.138 |
| **(25N,5.0cm)** | $F = 279.34$ $p < 0.001$* | *p*<0.001* | *p*<0.001* | *p*<0.001* | *p*<0.001* | *p*<0.001* | *p*<0.001* |
| **(25N,7.5cm)** | $F = 209.30$ $p < 0.001$* | *p*=0.049* | *p*<0.001* | *p*<0.001* | *p*<0.001* | *p*<0.001* | *p*<0.001* |
